# DNA damage-specific effects of Tel1/ATM and γH2A/γH2AX on checkpoint signaling in *Saccharomyces cerevisiae*

**DOI:** 10.1101/572271

**Authors:** Jasmine Siler, Na Guo, Zhengfeng Liu, Yuhua Qin, Xin Bi

## Abstract

DNA lesions trigger the activation of DNA damage checkpoints (DDCs) that stop cell cycle progression and promote DNA damage repair. *Saccharomyces cerevisiae* Tel1 is a homolog of mammalian ATM kinase that plays an auxiliary role in DDC signaling. γH2A, equivalent to γH2AX in mammals, is an early chromatin mark induced by DNA damage that is recognized by a group of DDC and DNA repair factors. We find that both Tel1 and γH2A negatively impact G2/M checkpoint in response to DNA topoisomerase I poison camptothecin independently of each other. γH2A also negatively regulates DDC induced by DNA alkylating agent methyl methanesulfonate. These results, together with prior findings demonstrating positive or no roles of Tel1 and γH2A in DDC in response to other DNA damaging agents such as phleomycin and ionizing radiation, suggest that Tel1 and γH2A have DNA damage-specific effects on DDC. We present data indicating that Tel1 acts in the same pathway as Mre11-Rad50-Xrs2 complex to suppress CPT induced DDC possibly by repairing topoisomerase I-DNA crosslink. On the other hand, we find evidence consistent with the notion that γH2A regulates DDC by mediating the competitive recruitment of DDC mediator Rad9 and DNA repair factor Rtt107 to sites of DNA damage. We propose that γH2A serves to create a dynamic balance between DDC and DNA repair that is influenced by the nature of DNA damage.

## 1. Introduction

Genotoxins cause genome instability by damaging DNA and/or blocking DNA replication. Cells have evolved intricate mechanisms for safeguarding genome integrity that are collectively called DNA damage response (1). DNA damage response recognizes DNA lesions, activates checkpoint pathways to arrest cell cycle progression, stabilizes DNA replication forks during S phase, and promotes DNA damage repair. In mammals, DNA damage checkpoint (DDC) signaling requires both the apical ATM and ATR kinases (2), whereas in the yeast *Saccharomyces cerevisiae*, the ATR homolog Mec1 plays an essential role in DDC, but the ATM homolog Tel1 only plays an auxiliary role that is usually masked by the prevailing activity of Mec1 (3–6).

The activation of DDC is intimately coupled with the generation of single stranded DNA (ssDNA) due to DSB end resection or replicative stress (7,8). The 3’ ssDNA resulted from DSB end resection also allows DSB repair by homology-based repair mechanisms. DSBs are recognized by Mre11-Rad50-Xrs2 (MRX) complex together with Sae2 (9) (Fig. 1B). MRX recruits Tel1 to DSBs and activates its kinase activity (9–11). Tel1 then supports MRX function in a positive feedback loop by stabilizing MRX association with DSBs, which is important for MRX mediated DSB repair (9,12). In addition, Tel1 phosphorylates Sae2, stimulating its activity (13,14) (Fig. 1B). Tel1 also phosphorylates histone H2A at its carboxyl-terminal serine 129 (equivalent to serine 139 of histone variant H2AX in mammals) creating H2A-S129-P, or γH2A (equivalent to mammalian γH2AX) containing nucleosomes (15) (Fig. 1B). Note H2A-S129 in chromatin surrounding DSB is also subject to phosphorylation by Mec1 later during DDC signaling (7,16–18) (Fig. 1D).

**Fig. 1.**
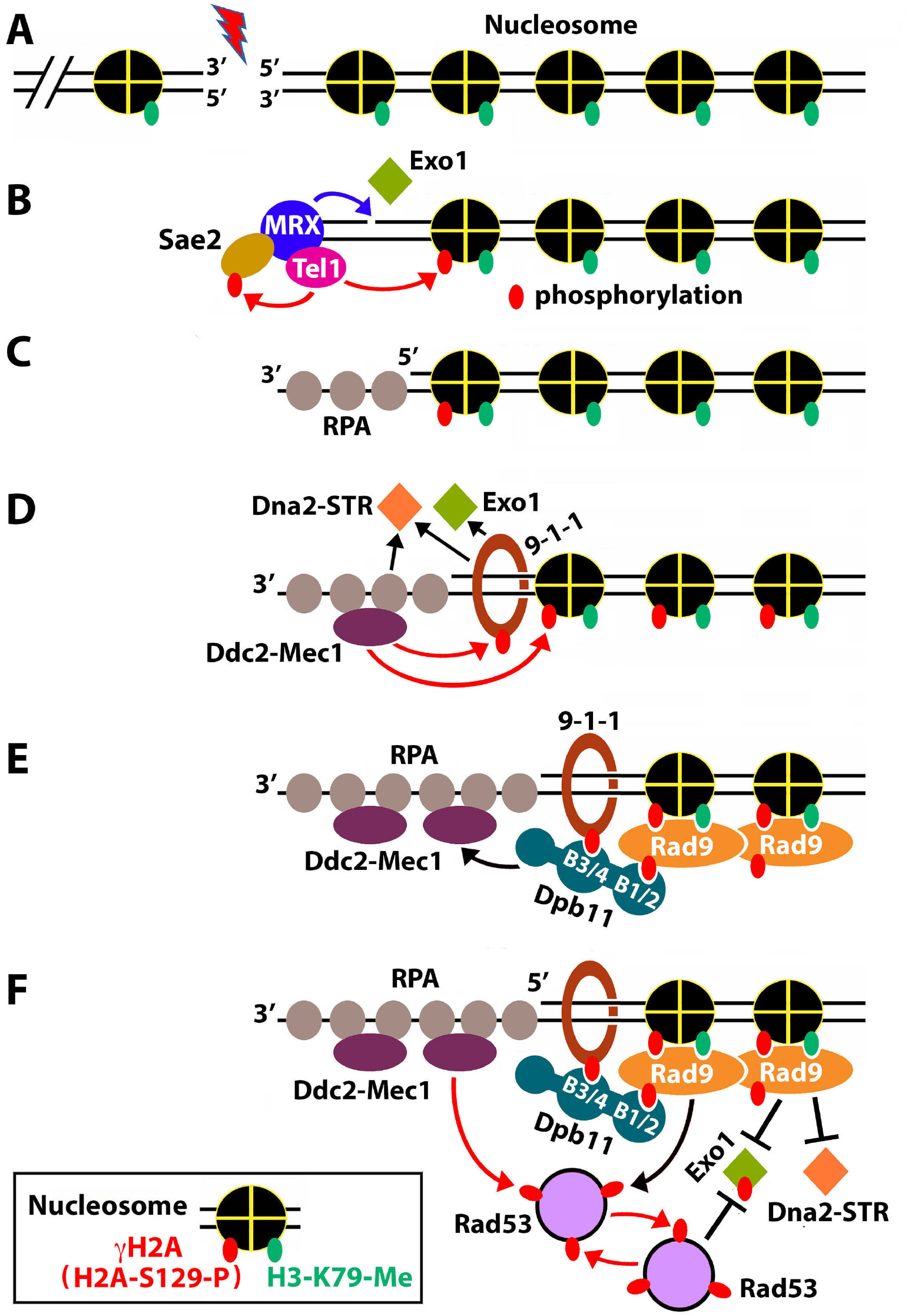
Mechanism of DNA damage checkpoint activation in *S. cerevisiae.* Red arrows denote phosphorylation. Black arrows denote recruitment and/or activation. Phosphorylation of Ser129 of histone H2A and methylation of lysine 79 of histone H3 on the nucleosome are indicated in the inset. See the text for descriptions.

The Mre11 subunit of MRX has both ssDNA endonuclease activity and 3’à5’ dsDNA exonuclease activity (19–22). Sae2 activates the ssDNA endonuclease activity of Mre11 that is responsible for the incision of the 5’ strand of DSB (23,24) (Fig. 1B, blue arrow). MRX also helps to recruit the 5’➔3’ exonuclease Exo1 and Dna2 endonuclease complexed with Sgs1 helicase, Top3 and Rmi1 (Dna2-STR) to DSBs (25). The combined action of exonuclease activities of Mre11 and Exo1 then carry out the initial resection of the 5’ strand of DSB, leading to a short 3’ ssDNA overhang covered by the ssDNA binding complex RPA (Fig. 1C). DNA damage clamp 9-1-1 (Rad17-Mec3-Ddc1) binds the junction between dsDNA and 5’ ssDNA (ds/ssDNA junction) (8) (Fig. 1D). 9-1-1 and ssDNA-RPA activate Exo1 and Dna2-STR, allowing them to perform more extensive (long range) DNA end resection (8) (Fig. 1D).

The generation of 3’ ssDNA triggers the dissociation of Tel1 from DSBs and termination of its signaling function, while promoting the activation of Mec1-dependent signaling (7,26). Mec1 (in complex with Ddc2) is recruited to DSBs by binding RPA associated ssDNA (7,8) (Fig. 1D). Mecl phosphorylates Ddcl of 9-1-1 and histone H2A (which maintains and expands γH2A containing chromatin) (7,16–18,27) (Fig. 1D). Phosphorylated 9-1-1 recruits the scaffold protein Dpb11 (human TopBP1 homolog) that performs multiple functions in initiation of DNA replication, DDC signaling, and DNA repair (28–30). Dpb11 functions in DDC by activating Mec1 and helping recruit DDC adaptor/mediator kinase Rad9 (28,31,32) (Fig. 1E). Dpb11 bears 4 BRCT domains (referred to as B1 to B4 here) that are known to interact with phosphoproteins. Dpb11 binds phosphorylated 9-1-1 via its BRCT motifs 3+4 (B3/4), and phosphorylated Rad9 via its B1/2 (Fig. 1E). Note, Rad9 can be phosphorylated by CDK1 in a cell cycle dependent fashion independently of DNA damage, and by Mec1 in response to DNA damage (7,33,34). Besides binding Dpb11, Rad9 also recognizes γH2A and methylated lysine 79 of histone H3 (H3-K79-me) via its double BRCT and double Tudor domains, respectively (35–37) (Fig. 1E). Note whereas the formation of γH2A is induced by DNA damage, H3-K79 methylation by Dot1 occurs independently of DNA damage (37).

Upon activation via Mec1-dependent phosphorylation, Rad9 binds and activates the checkpoint effector kinase Rad53 that then undergoes intermolecular autophosphorylation (33,38,39) (Fig. 1F). Rad53 is also phosphorylated by Mec1 (39,40) (Fig. 1F). Mec1 and Tel1 also phosphorylate and activate Chk1 kinase (not illustrated in Fig. 1) (41). Activated Rad53 and Chk1 molecules are then released from sites of DNA damage to transduce signals to downstream targets as part of a signaling cascade leading to cell cycle arrest and DNA repair (8). In the above-described DDC mechanism, ssDNA and ds/ssDNA junction resulted from DSB end resection are critically required for checkpoint signaling (Fig. 1D-F). Note that long (~200 nt) ssDNA gaps can also be generated at stalled replication forks during replicative stress, which would potentially also trigger Rad9-dependent DDC similarly as would ssDNA generated by DSB end resection (42,43) (Fig. 1C-F).

As the processes of DDC and DNA end resection are intertwined with each other, they may influence each other in multiples ways. For example, DDC negatively impacts DNA end resection. Exo1 activity is inhibited by its phosphorylation by Mec1 (44) (Fig. 1F). Moreover, Rad9 associated with chromatin is believed to serve as a physical barrier to DNA resection by Dna2-STR (45–48) (Fig. 1F). Recently, the Fun30 chromatin remodeler has been shown to promote long range DSB resection by counteracting Rad9 (49–51).

Activation of DDC and recovery from it need to be orchestrated to both allow time for DNA repair and ensure timely resumption of cell cycle progression afterwards. However, there is only limited understanding of mechanisms for the down regulation DDC that are important for preventing DDC hyperactivation and promoting DDC recovery. Protein phosphatases PP2C and PP4 have been found to dephosphorylate Rad53-P, thereby aiding in recovery from cell cycle checkpoints (52–54). Moreover, Sae2 and the Slx4/Rtt107 complex, a DNA repair scaffold, have been shown to negatively regulate DDC signaling (55,56). Sae2 limits the retention of MRX at DSBs, thereby preventing heightened DDC signaling by MRX if it is persistently associated with DSBs (55,57). Slx4/Rtt107 competes with Rad9 for recruitment to chromatin at sites of DNA damage, thereby reducing DDC signaling (29,56).

We recently found that Fun30 negatively impacts DDC signaling induced by camptothecin (CPT), a DNA topoisomerase I (Top1) poison that traps Top1 when it is crosslinked to DNA during its enzymatic function of relaxing DNA supercoiling (58). While investigating how Fun30 modulates DDC, we discovered that, interestingly, deleting Tel1 or blocking H2A-S129 phosphorylation (γH2A) enhanced CPT-induced phosphorylation of Rad53 and Rad9, which points to negative roles of Tel1 and γH2A in DDC. We showed that Tel1 and γH2A suppress G2/M checkpoint in response to CPT. Moreover, found that γH2A also negatively regulates DDC induced by DNA alkylating agent methyl methanesulfonate. We obtained evidence consistent with the hypothesis that Tel1 functions in the same pathway as MRX in processing DNA-protein crosslinks, whereas γH2A and Fun30 modulate the access of Rad9 to sites of DNA damage, thereby regulating DDC.

## 2. Materials and Methods

### 2.1. Yeast strains

Yeast strains used in this work are listed in Table 1. Gene deletion was done by replacing the ORF of the gene of interest with *KanMX, NatMX* or *TRP1*, which was verified by Southern blotting or PCR. Strains W303a, MC42-2d, RCY337-26a, RCY337-6c, and RCY337-3d were obtained from Dr. Thomas Petes (Duke University); W303-1A, SKY2939, QY364 and QY375 from Dr. Stephen Kron (University of Chicago); JKM139 and R726 from Dr. James Haber (Brandeis University); and YXC723 from Dr. Grzegorz Ira (Baylor College of Medicine).

**Table 1.**
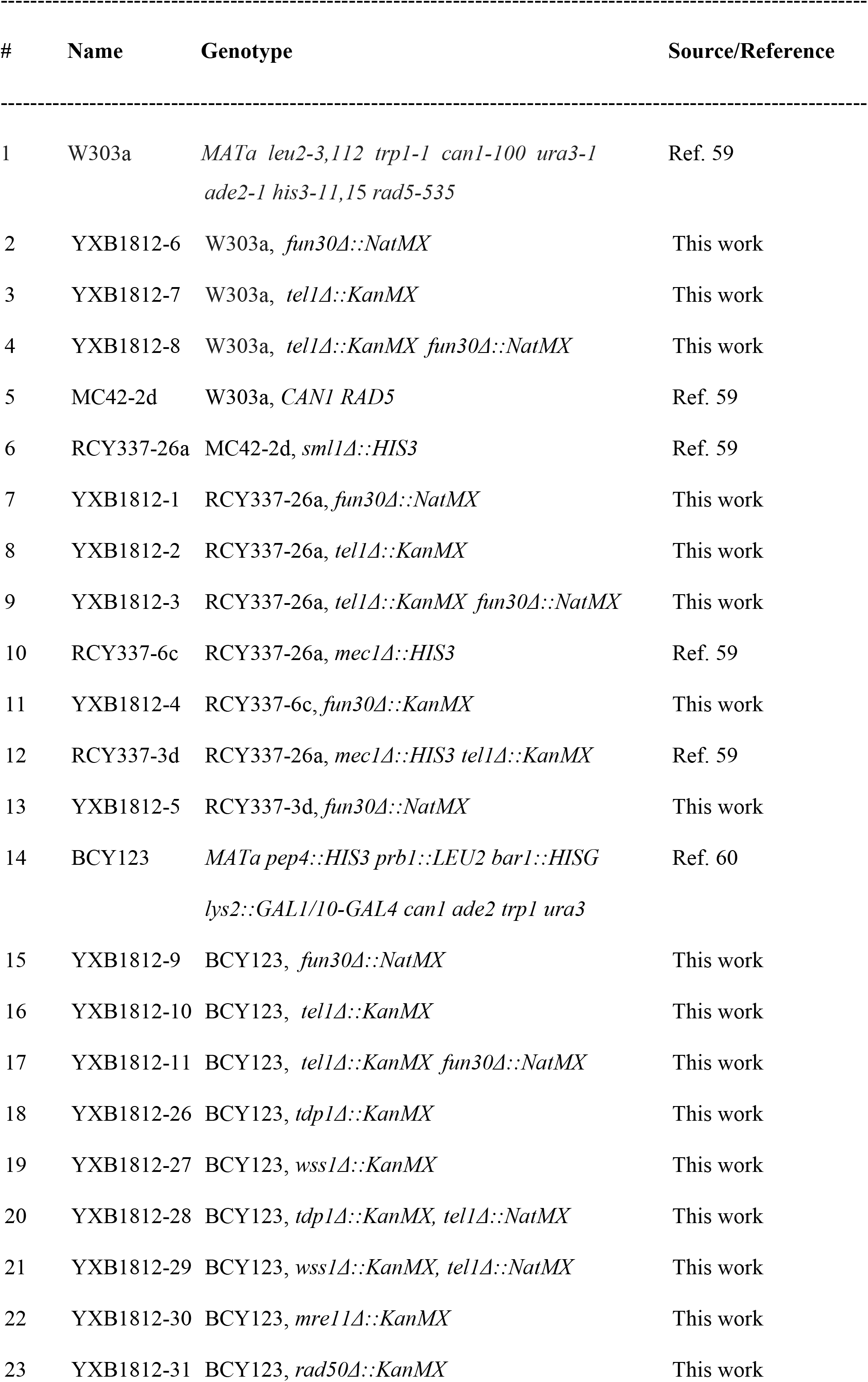
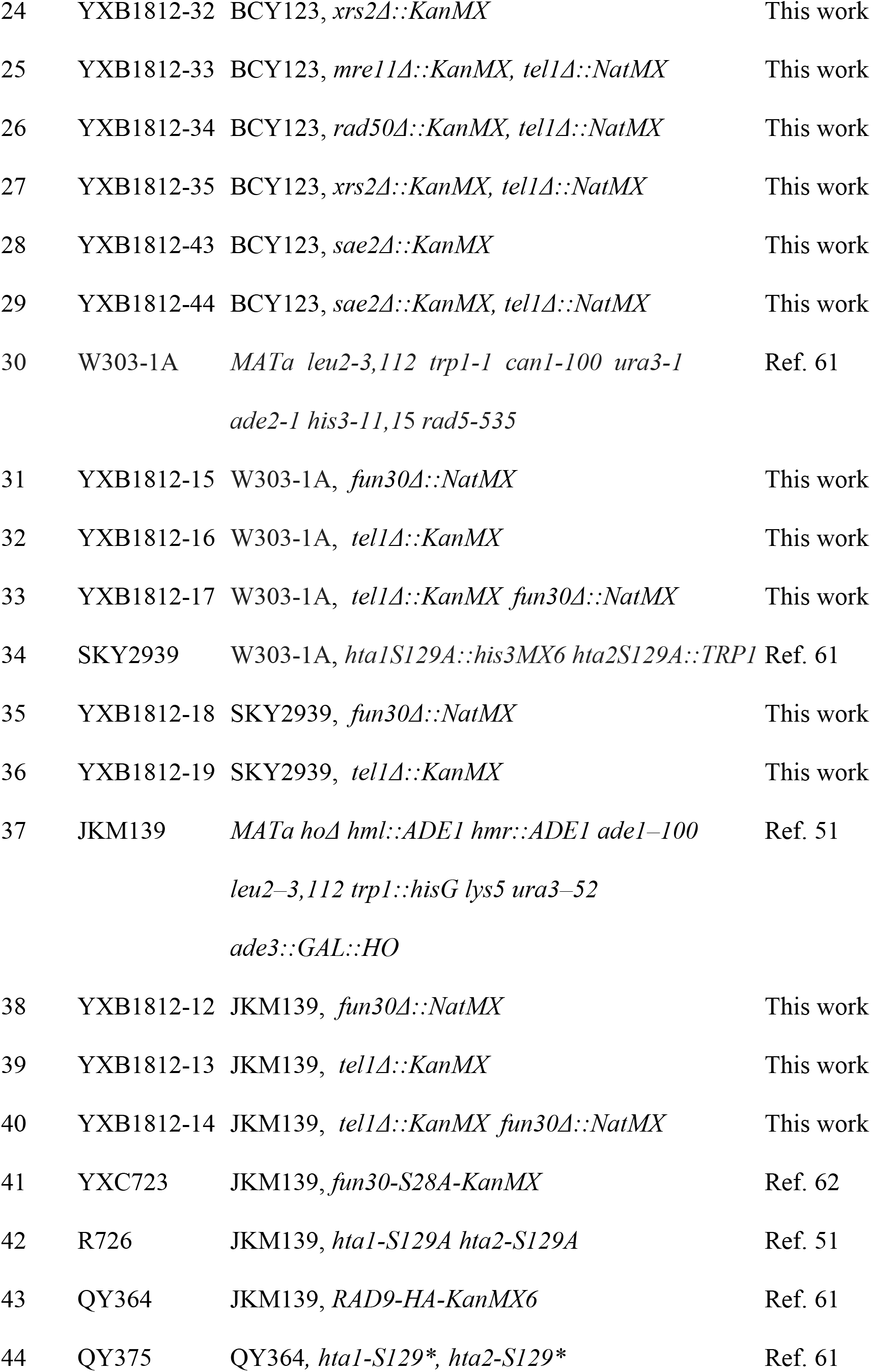
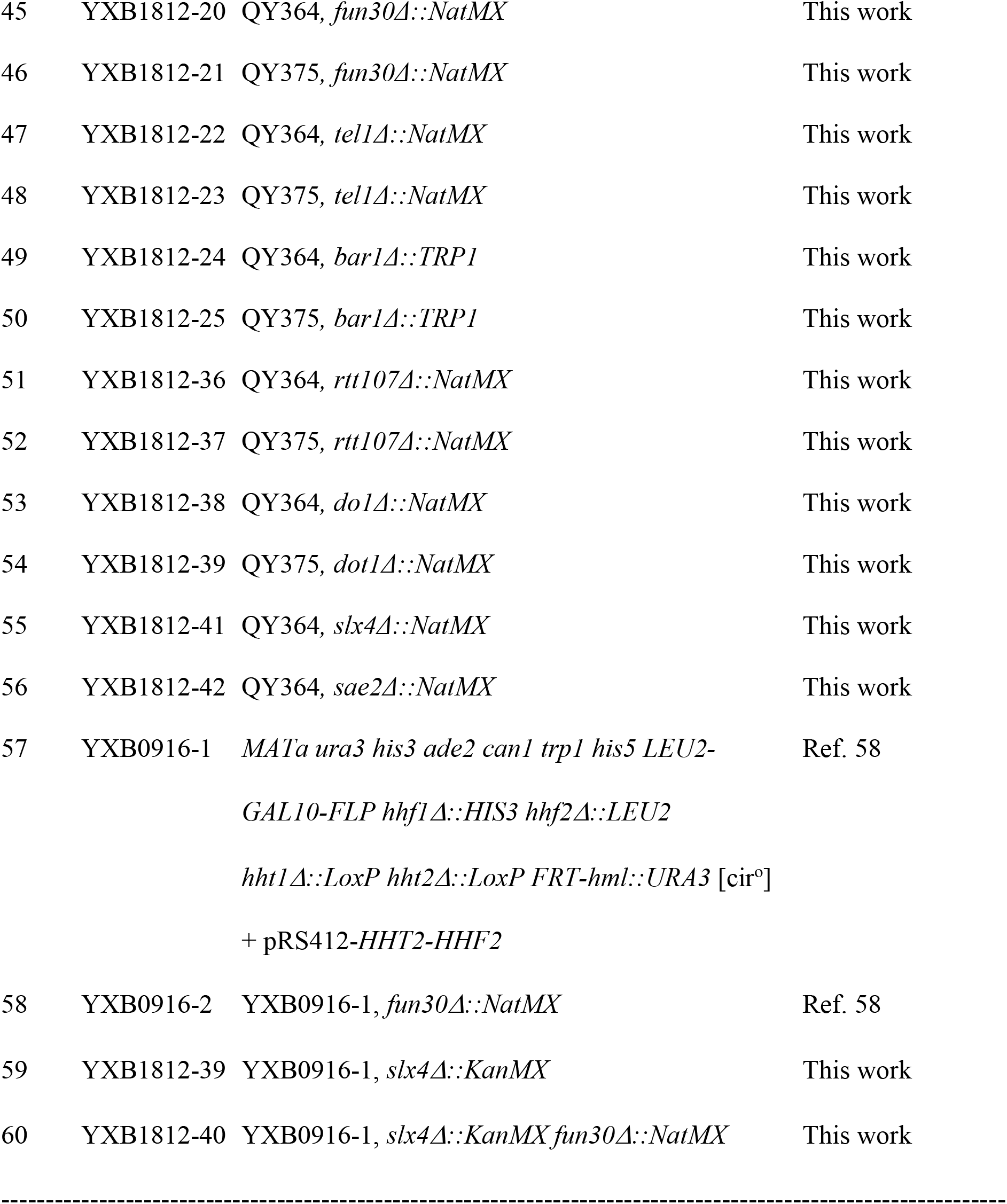
Yeast strains

### 2.2. SDS-PAGE and Western blotting

Proteins were isolated from yeast by TCA extraction as described (63). Ten μg of proteins from each sample of cells was subjected to SDS-PAGE in 4-12% gradient gels. Western blotting was performed using LI-COR Odyssey CLx Infrared Imaging System (LI-COR Biosciences) (63). Antibodies used in the work were: goat polyclonal anti-Rad53 (yC-19: sc6749, Santa Cruz Biotechnology), rabbit polyclonal anti-Clb2 (y-180: sc9071, Santa Cruz Biotechnology), rabbit polyclonal anti-HA (H6908, Sigma-Aldrich), rabbit polyclonal anti-G6PD (A9521, Sigma-Aldrich). Secondary antibodies used are LI-COR IRDye 800CW goat polyclonal anti-rabbit IgG (H+L) 926-32211 and LI-COR IRDye 800CW donkey anti-goat IgG (H+L) 926-32214.

### 2.3. Fluorescence activated cell sorting (FACS)

FACS analyses of yeast cells were performed as described (63) and analyzed on a FACSCalibur (Becton, Dickinson and Company). Data analysis was done using FlowJo software.

## 3. Results

### 3.1. Tel1 negatively impacts G2/M DNA damage checkpoint in response to CPT independently of Fun30

We recently found that Fun30 has a negative effect on Rad53 phosphorylation (Rad53-P) induced by CPT (58). Since Rad53 can be phosphorylated by both Mec1 and Tel1, we wondered if Fun30 compromises the abilities of Mec1 and/or Tel1 to phosphorylate Rad53. To address this question, we examined the level of Rad53-P in CPT-treated *fun30Δ* and wild type (WT) cells lacking Mec1 and Tel1 individually or simultaneously. Rad53 can be phosphorylated at multiple serine or threonine (S/T) residues (39,64), and Rad53-P molecules migrate more slowly than Rad53 in SDS-PAGE as a smear and/or band(s) above the band corresponding to Rad53. As such, the level of Rad53-P in a particular culture of cells can be estimated as the intensity of Rad53-P species relative to that of Rad53 on a Western blot. As shown in Fig. 2A, there was a moderate level of Rad53-P in exponentially growing WT cells treated with 5 μg/ml (14 μM) of CPT (compare lane 1c with 1). This is consistent with the previous finding that CPT triggers only a modest checkpoint response, despite its ability to slow down DNA replication forks and potentially induce DSBs (58,65–67). Rad53-P abundance in *fun30Δ* cells was significantly higher than that in WT cells (Fig. 2A, compare lane 2c with 1c), confirming our prior observation (58). Little or no CPT-induced Rad53-P was present in *mec1Δ* and *mec1Δ fun30Δ* mutants (Fig. 2A, lanes 5c and 6c). On the other hand, Rad53-P still existed in *tel1Δ* cells, and its abundance was increased by *fun30Δ* (Fig. 2A, lanes 3c with 4c). These results demonstrate that it is Mec1-mediated, not Tel1-mediated, Rad53-P in response to CPT that is negatively impacted by *fun30Δ.*

**Fig. 2.**
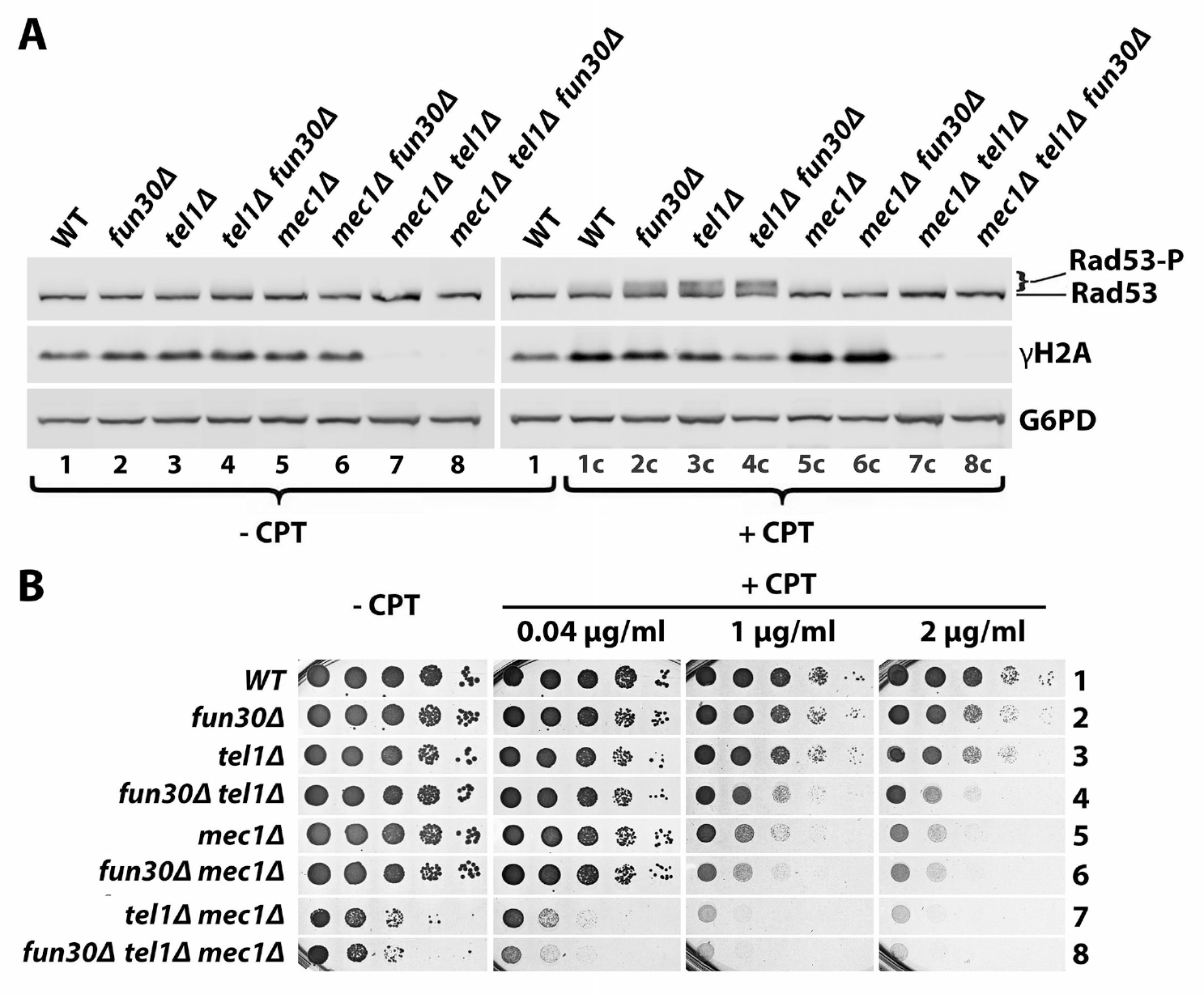
Tel1 negatively impacts DDC and promotes CTP resistance independently of Fun30. (A) Western blot analysis of Rad53, γH2A and G6PD from indicated strains (listed as #6-13 in Table 1) with or without CTP treatment. Exponentially growing cells were treated with (+CPT) or without (-CPT) CPT at 5 μg/ml for 90 minutes. Protein extracts form these cells were subjected to SDS-PAGE and Western blotting followed by detection of Rad53, γH2A and G6PD (glucose-6-phosphate dehydrogenase) with respective antibodies. (B) Growth phenotypes of indicated strains (#6-13 in Table 1) on media with or without CTP. Ten-fold serial dilutions of a late log phase culture of each strain were plated on synthetic complete plates with or without CPT, and incubated for three days.

Notably, Rad53-P in CPT-treated *tel1Δ* cells was robust and clearly more abundant than that in WT cells (Fig. 2A, compare 3c with 1c). Therefore, Tel1 *per se* plays a negative role in Rad53-P in response to CPT, which is apparently at odds with the notion that Tel1 serves as an auxiliary DDC activator (3,4,8). We wondered whether this unexpected phenotype of *tel1Δ* was specific to the genetic background of the cells used in our experiment. Our strains were derived from RCY337-26a bearing the *sml1Δ* mutation that offsets the lethal effect of *mec1Δ* mutation (Table 1) (59). Since Sml1 is involved in DNA damage response, it is possible that the effect of *tel1Δ* on Rad53-P observed was dependent on *sml1Δ.* To test this notion, we repeated the above experiment using strains derived from W303-1A, BCY123, or JKM139 (all of which contain intact *SML1)* (51,60,61) (Table 1) and obtained the same result, namely *tel1Δ* increased CPT-induced Rad53-P (Fig. S1, A-C). Moreover, Menin et al. also reported recently a similar finding (68) while this manuscript was under preparation. Therefore, Tel1 appears to exert a negative impact on DDC signaling in response to CPT. The fact that Rad53-P in *tel1Δ fun30Δ* double mutant was more robust than that in *tel1Δ* and *fun30Δ* single mutants and WT strain (Fig. 2A and S1) suggests that Tel1 and Fun30 function independently of each other to hinder CPT-induced DDC signaling.

In the presence of DNA damage, checkpoint pathways may be activated in G1, S and/or G2/M phases of the cell cycle (1,8). Given that Fun30 and Tel1 negatively impact CPT-induced Rad53-P in cycling cells (Fig. 2A), we wondered whether they promote cell cycle progression in the presence of CPT by countering S and/or G2/M checkpoints. To address this question, we monitored the synchronous progression of *fun30Δ* and *tel1Δ* single and double mutants as well as WT cells in the cell cycle in the presence of CPT following their release from G1 arrest. Asynchronous cells (designated Asy) were first arrested in G1 by α-factor and then released into fresh medium with or without 5 μg/ml CPT for a 105 minute incubation. Aliquots of the culture were taken every 15 minutes for monitoring Clb2 and Rad53-P in them as indicators of cell cycle progression and checkpoint signaling, respectively. Clb2 is a B-type cyclin involved in the control of G2/M transition that steadily accumulates during G2 and disappears at mitosis (69). Mitochondrial porin (Por1) was also detected as a loading control.

We found that in the absence of CPT, Clb2 in WT cells was not detectable in G1 and 15 min after G1 release, but started to appear 30 min after G1 release, peaked around 45 min and subsided afterwards (Fig. 3, row 1 in Clb2 panel). This indicates that WT cells entered G2 around 30-45 min and exited mitosis around 75-90 min. Similar results were obtained for *fun30Δ* mutant (Fig. 3, compare row 2 to 1 in Clb2 panel), suggesting that Fun30 is not required for cells to traverse S and G2 phase or exit from mitosis. Tel1 also did not affect S phase progression, but slightly delayed the exit from mitosis (Fig. 3, compare row 3 to 1 in Clb2 panel). Consistent with that neither *fun30Δ* nor *tel1Δ* caused a significant delay in cell cycle progression, Rad53-P was not readily detectable in *fun30Δ* or *tel1Δ* cells after G1 release (Fig. 3, rows 2 and 3 in Rad53 panel). The *tel1Δ fun30Δ* double mutation had little or no effect on S or G2 progression, but moderately delayed the exit from mitosis (Fig. 3, row 4 in Rad53 panel). This was accompanied by a modest accumulation of Rad53-P at 75 min and later time points after G1 release (Fig. 3, compare row 4 to 1 in Rad53 panel). Taking together, the above results suggest that Fun30 and Tel1 have small and redundant roles in promoting efficient G2/M transition during unperturbed cell growth.

**Fig. 3.**
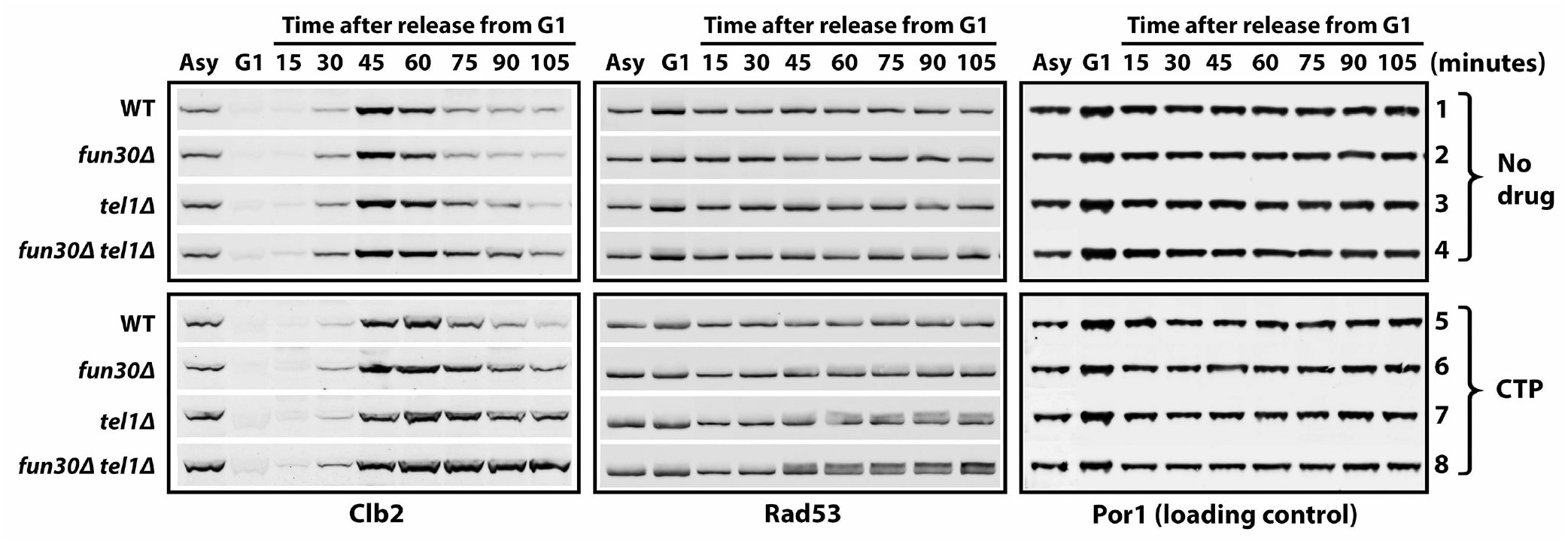
Tel1 and Fun30 down regulate CPT-induced G2/M checkpoint independently of each other. Asynchronous (Asy) cells of the indicated strains (#14-17 in Table 1) were arrested in G1 by a 2-hour α-factor treatment, and then released into medium with or without (No drug) 5 μg/ml CTP, and incubated for 105 minutes. Aliquots of cells were collected at the indicated time points during CTP treatment. Clb2, Rad53 and Por1 in protein extracts from the aliquots were examined by SDS-PAGE and Western blotting.

CPT caused little or no delay in S phase progression of WT cells (Fig. 3, compare rows 1 and 5 in Clb2 panel), which is in line with the notion that CPT does not induce a robust intra-S checkpoint (58,65). In fact, CPT-induced DNA damage in S phase was previously considered checkpoint blind (65). We found that none of *fun30Δ* and *tel1Δ* single and double mutants had any significant delay in S phase progression in the presence of CPT (Fig. 3, Clb2 panel, compare rows 6-8 with 2-4, respectively, at 15, 30 and 45 min). This suggests that the apparent lack of an intra S-phase checkpoint in the presence of CPT is not because the checkpoint is suppressed by Fun30 or Tel1. On the other hand, we observed that *fun30Δ, tel1Δ* and *fun30Δ tel1Δ* mutants experienced an increasingly more severe retardation in the exit from mitosis, as reflected by the delayed disappearance of Clb2 after 45 min of incubation with CPT (Fig. 3, Clb2 panel, compare rows 6-8 with 5), which was correlated with increasingly higher levels of Rad53-P found in these strains (Fig. 3, Rad53 panel, compare rows 6-8 with 5). These results indicate that CPT induces a G2/M arrest that is modest in *fun30Δ* cells, robust in *tel1Δ* cells and strongest in *fun30Δ tel1Δ* cells. Therefore, we conclude that Fun30 and Tel1 independently mitigate a G2/M checkpoint induced by CTP.

Since a loss in proper regulation of DDC may result in a decline in cell survival in the presence of a genotoxin, we examined the effects of *tel1Δ, fun30Δ* and *mec1Δ* single, double and triple mutations on cell survival in the presence of CPT. As shown in Fig. 2B and S1A-C, each double mutant was more sensitive to CPT than the corresponding single mutants, and the triple mutant more sensitive to double and single mutants. This indicates that Tel1, Fun30 and Mec1 can contribute to CPT-resistance independently of each other. It is noteworthy that *heightened*

DDC signaling in *tellΔ* and *fun30Δ* single and double mutants in response to CPT (Fig. 2A and S1A-C) was correlated with *reduced* CPT-resistance of these mutants (Fig. 2B and S1A-C). On the other hand, *diminished* DDC activation in *mec1Δ* mutant (Fig. 2A) was also correlated with *reduced* CPT-resistance (Fig. 2B). Therefore, there does not appear to be a simple, straightforward correlation between the level of DDC signaling and CPT-resistance.

### 3.2. γH2A also plays a negative role in CPT-induced G2/M checkpoint

Histone H2A-S129 phosphorylation, or γH2A (equivalent to γH2AX in mammals), is an early chromatin mark in response to DNA damage or DNA replication stress (1,16–18) (Fig. 1). Phosphorylation of H2AX/H2A is carried out redundantly by ATM/Tel1 and ATR/Mec1 kinases (1,17,70). Consistently, we showed that γH2A was there in cells deleted for Mec1 or Tel1, but was abolished in cells lacking both Mec1 and Tel1 (Fig. 2A, 3, 5, 7, 3c, 5c and 7c in γH2A panel). We wondered whether the effect of Tel1 on CPT-induced Rad53-P was related to its role in phosphorylating histone H2A. In addition, given that γH2A has been shown to hinder Fun30 binding to nucleosomes *in vitro* (51), we also wondered if γH2A influenced how Fun30 impacts CPT-induced Rad53-P. To address these questions, we measured Rad53-P in a series of *tel1Δ, fun30Δ*, and *hta-S129A* (with S129 of H2A changed to alanine) single and double mutants as well as WT cells upon CPT treatment.

We found that CPT-induced Rad53-P level in *tel1Δ hta-S129A* double mutant was higher than that in *hta-S129A* mutant (Fig. 4A). Rad53-P abundance in *fun30Δ hta-S129A* double mutant was similar with that in *hta-S129A* mutant (Fig. 4A). We corroborated these results by showing that Rad53-P in *tel1Δ hta-S129\** mutant was higher than that in *hta-S129\** mutant, whereas *fun30Δ hta-S129\** double mutant and *hta-S129\** mutant had similar levels of Rad53-P in the presence of CPT (Fig. S2A). Note H2A-S129* is a truncated allele of histone H2A deleted for its C-terminal four amino acids including S129, and is thus unable to be phosphorylated at S129 (16). The above results indicate that Tel1 modulates DDC independently of γH2A, whereas Fun30 does so in a γH2A-dependent fashion.

**Fig. 4.**
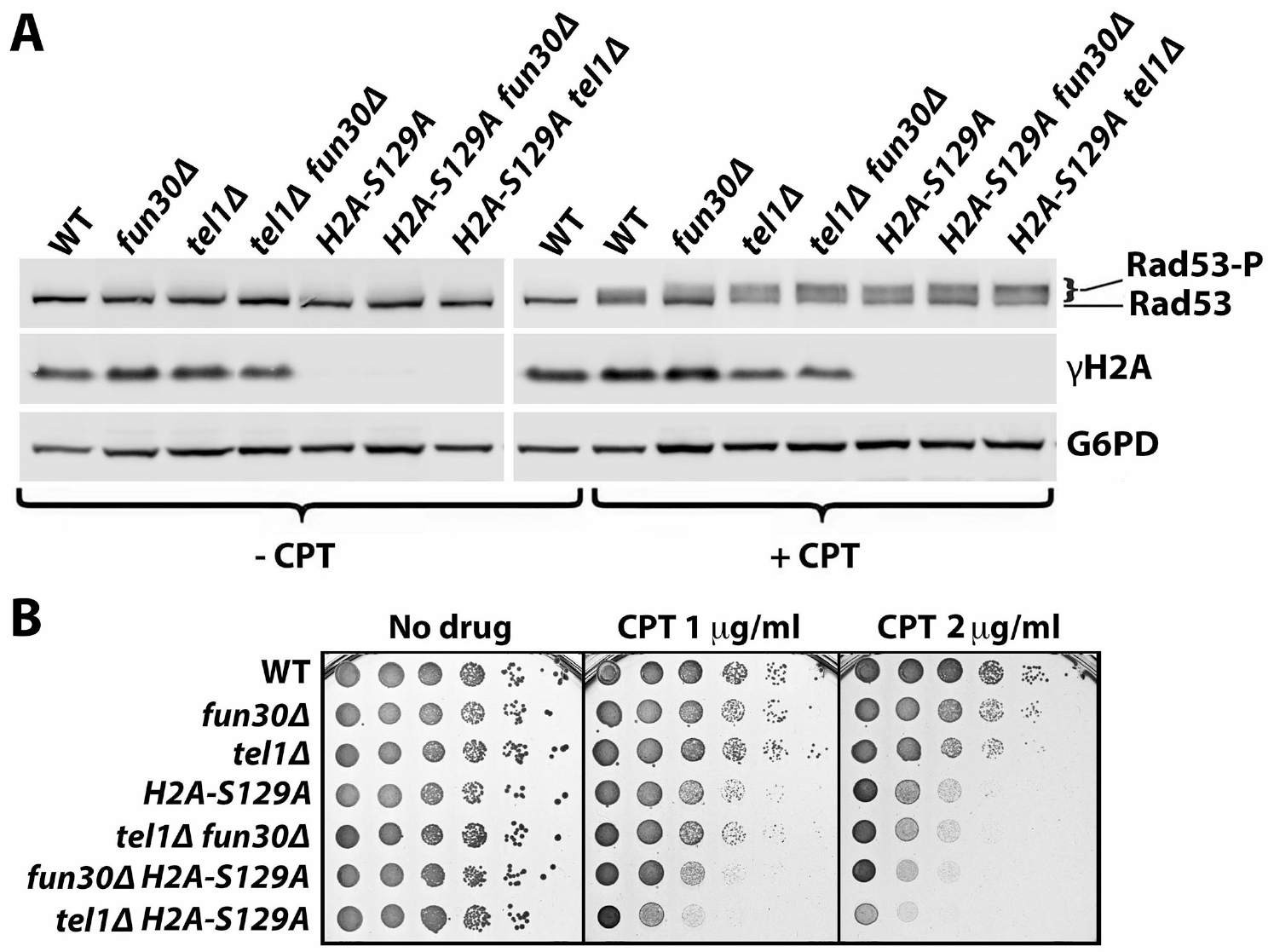
γH2A inhibits DDC signaling and promotes CTP-resistance independently of Tel1 and Fun30. (A) Western blot analysis of Rad53, γH2A and G6PD from indicated strains (#30-36 in Table 1) with (+CPT) or without (-CPT) CTP treatment. (B) Growth phenotypes of indicated strains (#30-36 in Table 1) on media with or without CTP.

Interestingly, we observed that Rad53-P in *hta-S129A* or *hta-S129\** mutant was clearly higher than that in wild type cells (Fig. 4A and S2A). In fact, *hta-S129A* or *hta-S129\** enhanced Rad53-P to a larger extent than *fun30Δ* did (Fig. 4A and S2A). These results demonstrate that γH2A also plays a negative role in DDC signaling in response to CPT. This role of γH2A is not dependent on Tel1 or Fun30 as *hta-S129A* or *hta-S129\** mutation increased Rad53-P in cells lacking Tel1 or Fun30 (Fig. 4A and S2A). CPT-sensitivity test of *hta-S129A, tel1Δ* and *fun30Δ* single and double mutants revealed that γH2A plays a larger role than Tel1 and Fun30 in CPT-resistance that is independent of Tel1 and Fun30 (Fig. 4B).

We next examined whether the increase in CPT-induced Rad53-P due to the lack of γH2A in cycling cells reflected an enhancement of DDC in S and/or G2/M phases. We arrested *hta-S129\** and WT cells in G1 by α-factor and then released them into fresh medium with or without CPT for a further 150 min incubation. Aliquots of the culture were taken every 15 minutes after G1 release for FACS and Western blotting analyses. As shown in Fig. 5A, FACS data showed that WT cells reached G2 about 45 min after G1 release, and started to exit mitosis at about 75 min in the absence of CPT (-CPT panel). At 150 min, the culture consisted of significant proportions of cells in G1, S and G2/M phases, which is similar to an asynchronous cell culture (Fig. 5A), likely reflecting a loss of synchrony in cell cycle progression after prolonged incubation. The presence of CPT had no effect on the progression of WT cells from G1 to G2 (up to ~60 min) (Fig. 5A, WT, compare-CPT and +CPT panels), demonstrating again that CPT does not induce an intra-S checkpoint. On the other hand, only a small portion of cells was able to exit mitosis in the presence of CPT (Fig. 5A, WT, 90-150 min), indicating again that CPT triggered a G2/M checkpoint response. The progression of *hta-S129\** cells in the cell cycle following G1 release in the absence of CPT was similar to that of WT cells (Fig. 5A, compare WT and *hta-S129\**,-CPT panels), suggesting that *hta-S129\** does not significantly affect cell proliferation. In the presence of CPT, *hta-S129\** did not delay S phase progression (Fig. 5A, compare WT+CPT, and *hta-S129*+CPT* panels, 15-60 min), but caused a tighter blockage of the exit from mitosis (Fig. 5A, compare WT+CPT, and *hta-S129*+CPT* panels, 75-150 min). These results indicate that *hta-S129\** enhances CPT-induced G2/M checkpoint.

**Fig. 5.**
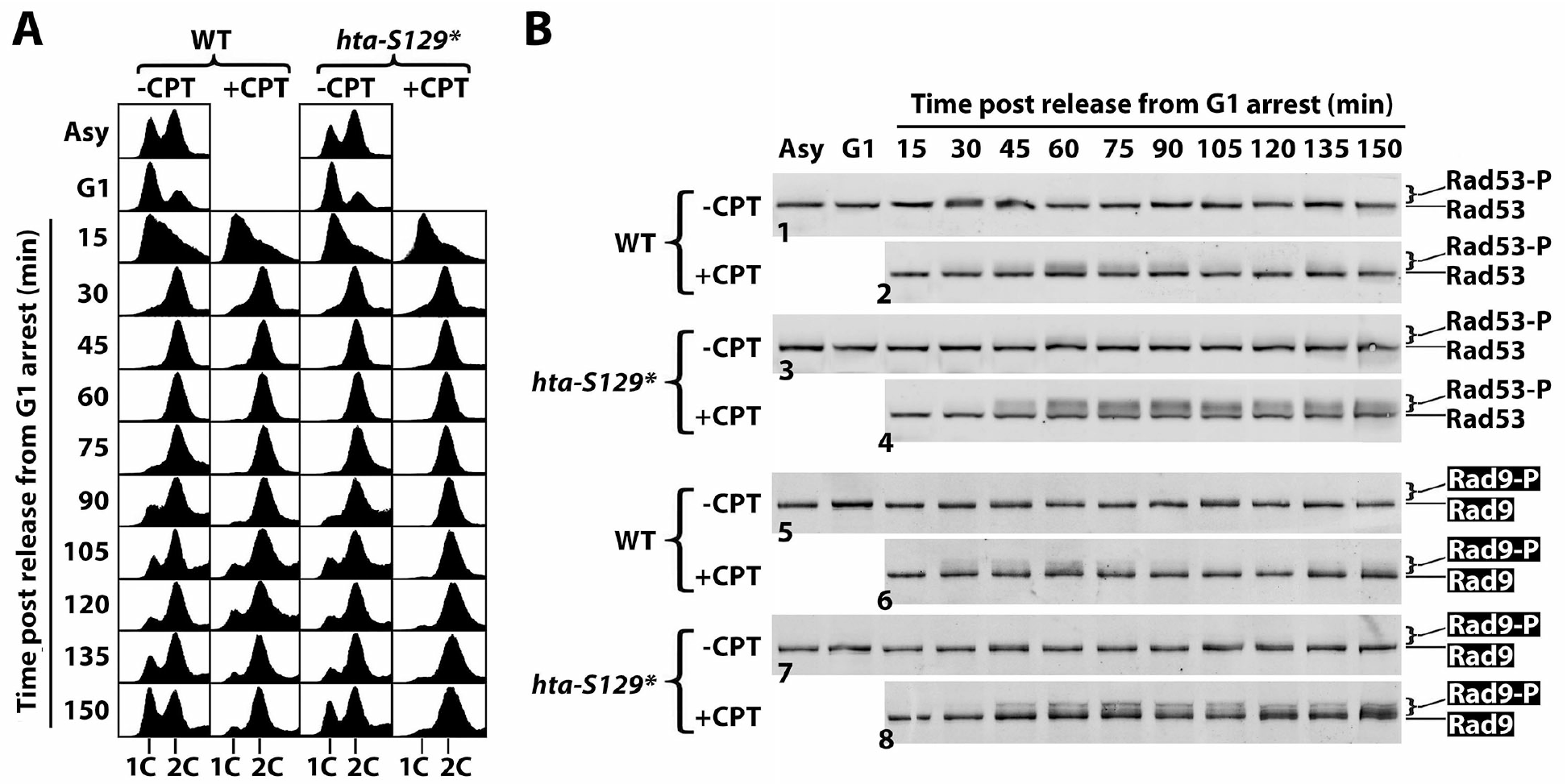
γH2A hinders G2/M checkpoint in response to CPT. Asynchronous (Asy) cells of WT (YXB1812-24) and *hta-S129\** (YXB1812-25) (#49 and 50 in Table 1) were arrested in G1 by α-factor (α-F), and released into medium with (+CPT) or without (-CPT) 5 μg/ml CTP, and incubated for 150 minutes. Aliquots of cells were collected at the indicated time points for FACS analysis (A) and protein extraction for examining Rad53 and HA-tagged Rad9 by SDS-PAGE and Western blotting (B).

Consistent with the above FACS results showing an enhancement of CPT-induced G2/M cell cycle checkpoint by *hta-S129\** mutation, we found that *hta-S129\** increased the level of CPT-induced Rad53-P when cells entered G2 phase (Fig. 5B, compare panels 2 and 4). This was accompanied by an increase in the phosphorylation of DDC adaptor Rad9 (Rad9-P) (Fig. 5B, compare panels 6 and 8). Therefore, γH2A negatively impacts the activation of CPT-induced DDC signaling. It is noteworthy that in WT cells, CPT-induced Rad53-P and Rad9-P peaked around 60 min when WT cells entered G2/M and declined afterwards (Fig. 5B, panels 2 and 6). On the other hand, in *hta-S129\** cells, Rad53-P and Rad9-P levels stayed high after reaching their maximum values between 60 and 75 min (Fig. 5B, panels 4 and 8). These results suggest that γH2A also negatively regulates the recovery of G2/M checkpoint.

### 3.3. Tel1 suppresses CPT-induced DDC in the same pathway as MRX, and independently of Tdp1 and Wss1

It is intriguing that Tel1 negatively impacts CPT-induced DDC (Fig. 2 and 3), while having positive or no effects on DDC triggered by other DNA damages including those inflicted by phleomycin or methyl methanesulfonate (MMS) as well as DSBs made by HO or restriction endonucleases (6,68,71,72). These findings suggest that Tel1 modulates DDC in a DNA damage-dependent manner. Phleomycin is a radiomimetic agent that generates single or double stranded DNA termini (73,74), and MMS alkylates DNA mainly on guanine and adenine (75). Note that these DNA lesions and DSBs made by endonucleases are protein-free, but CPT traps a unique crosslinked DNA-protein complex (DPC) called Top1 cleavable complex (Top1cc) consisting of a nick with its 3’ end covalently linked to Top1 via a phosphodiester bond (67,76). DPCs are processed/repaired differently from other types of DNA lesions. Specifically, DPCs can be repaired by three distinct mechanisms mediated by Tdp1 tyrosyl-DNA phosphodiesterase, Wss1 protease, and MRX nuclease, respectively, that target different parts of a DPC (76). Tdp1 hydrolyzes the covalent tyrosyl-DNA phosphodiester bond between Top1 and DNA, Wss1 degrades the protein moiety of DPC, whereas MRX attacks the DNA (76).

It is possible that Tel1 specifically participates in, or regulates, the processing of Top1cc, in a way that prevents the generation of DDC inducing DNA structures. We tested this hypothesis by examining whether Tel1’s effect on CPT-induced Rad53-P was dependent on any of the aforementioned factors involved in DPC repair (76). As shown in Fig. 6A, *tel1Δ* and *tdp1Δ* had a synthetic effect on CPT-induced Rad53-P, and so did *tel1Δ* and *wss1Δ* (Fig. 6A), indicating that Tel1 can negatively impact CPT-induced DDC signaling independently of Wss1 and Tdp1. Notably, we observed that similar to *tel1Δ, tdp1Δ* or *wss1Δ* alone also increased CPT-induced Rad53-P (Fig. 6A, +CPT panel). Moreover, *mre11Δ, rad50Δ* or *xrs2Δ* alone also elevated the level of CPT-induced Rad53-P (Fig. 6B, +CPT panel). These results suggest that the repair of Top1cc by Tdp1, Wss1 and MRX down regulates DDC signaling. Importantly, we found that deletion of Tel1 from *mre1Δ, rad50Δ* or *xrs2Δ* mutant didn’t affect the level of CPT-induced Rad53-P (Fig. 6B, +CPT panel), indicating that the negative impact of Tel1 on DDC is dependent on MRX. Therefore, Tel1 negatively impacts CPT-induced DDC in the same pathway as MRX, and independently of Tdp1 and Wss1.

**Fig. 6.**
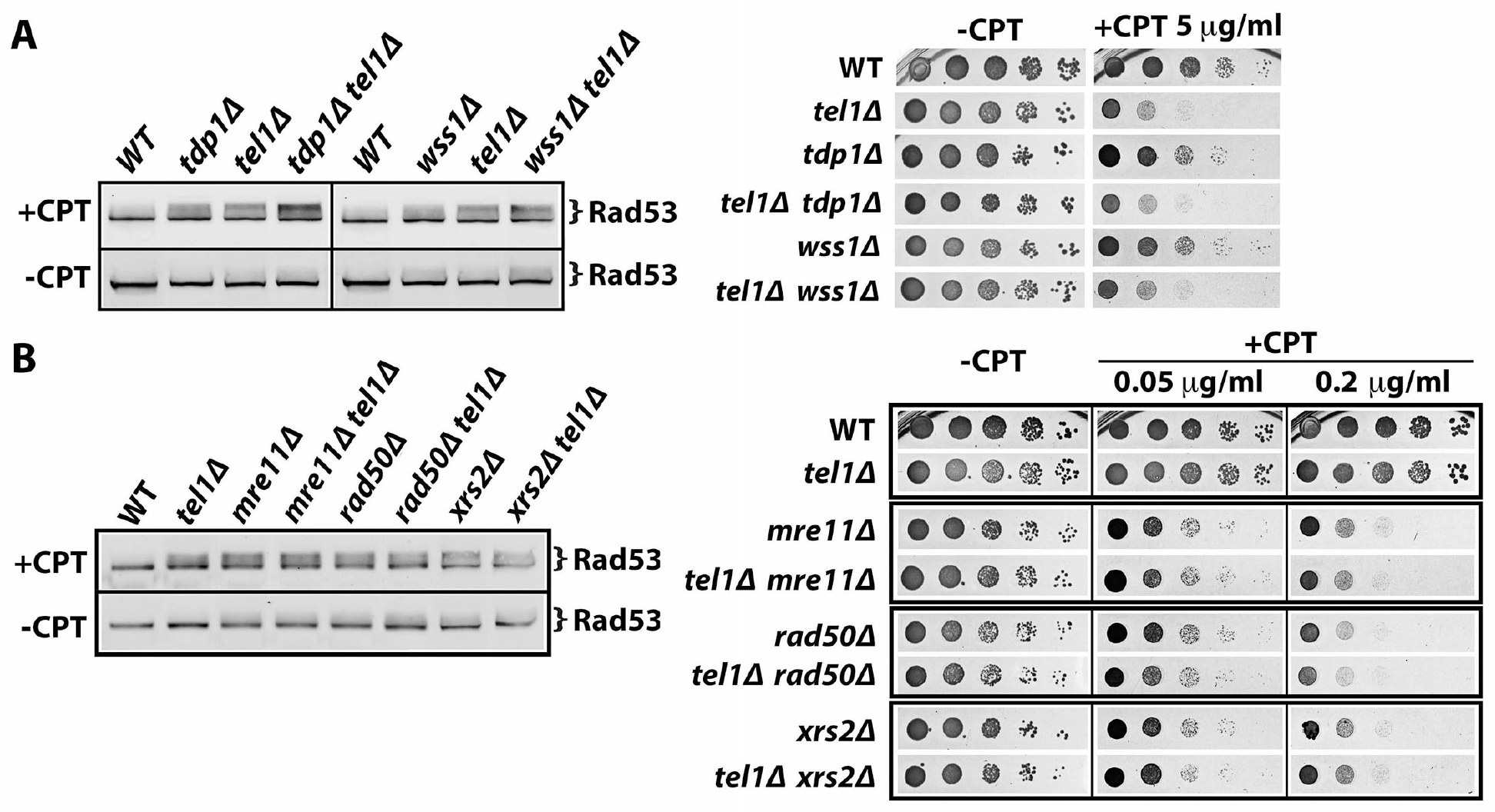
Tel1 negatively impacts CPT-induced DDC signaling in the same pathway as MRX and independently of Tdp1 and Wss1. Shown are results from Western blot analysis of Rad53 from indicated strains with or without CPT treatment as well as growth phenotypes of the strains on media with or without CTP. (A) Strains #14, 16 and 18-21 were tested. (B) Strains #14 and 16 and 22-27 were tested.

### 3.4. γH2A and Fun30 may down regulate DDC by hindering Rad9 recruitment to DNA lesions

Our finding that γH2A negatively impacts CPT-induced DDC (Fig. 4 and 5) is apparently counterintuitive as γH2A is known to help recruit Rad9 to chromatin at DNA lesions (35–37). This conundrum might stem from the competition between Rad9 and DNA repair scaffold Slx4/Rtt107 for binding γH2A and Dpb11 at DNA damage sites (29,56,77–79). Rad9 bears a double BRCT domain that recognizes γH2A, and Rad9 phosphorylated at S464 and T474 by CDK binds to the B1/2 domain of Dpb11 tethered to 9-1-1 complex (35,79) (Fig. 7A, left). On the other hand, the BRCT domain of Rtt107 also recognizes γH2A, and Slx4 phosphorylated by CDK also binds to B1/2 domain of Dpb11 (77,80) (Fig. 7A, right). The competition between Rad9 and Slx4/Rtt107 for binding γH2A and Dpb11 is believed to yield a dynamic balance between DNA lesions associated with Rad9 and those with Slx4/Rtt107 (56) (Fig. 7A). In line with this model, deletion of Rtt107 has been shown to increase DDC signaling likely by allowing Rad9 to bind γH2A and Dpb11 more efficiently (56) (Fig. S3B). Loss of γH2A (as in *hta-S129\** mutant) would eliminate γH2A-mediated recruitment of Rad9 and Slx4/Rtt107 to damaged chromatin, but Rad9 and Slx4/Rtt107 can still interact with Dpb11, and Rad9 can additionally associate with H3-K79-me (Fig. S3C). We propose that under this circumstance in *hta-S129\** mutant, Rad9 outcompeted Slx4/Rtt107 for engaging DNA lesions (Fig. S3C), resulting in an increase in DDC signaling as was shown in Fig. 4A.

**Fig. 7.**
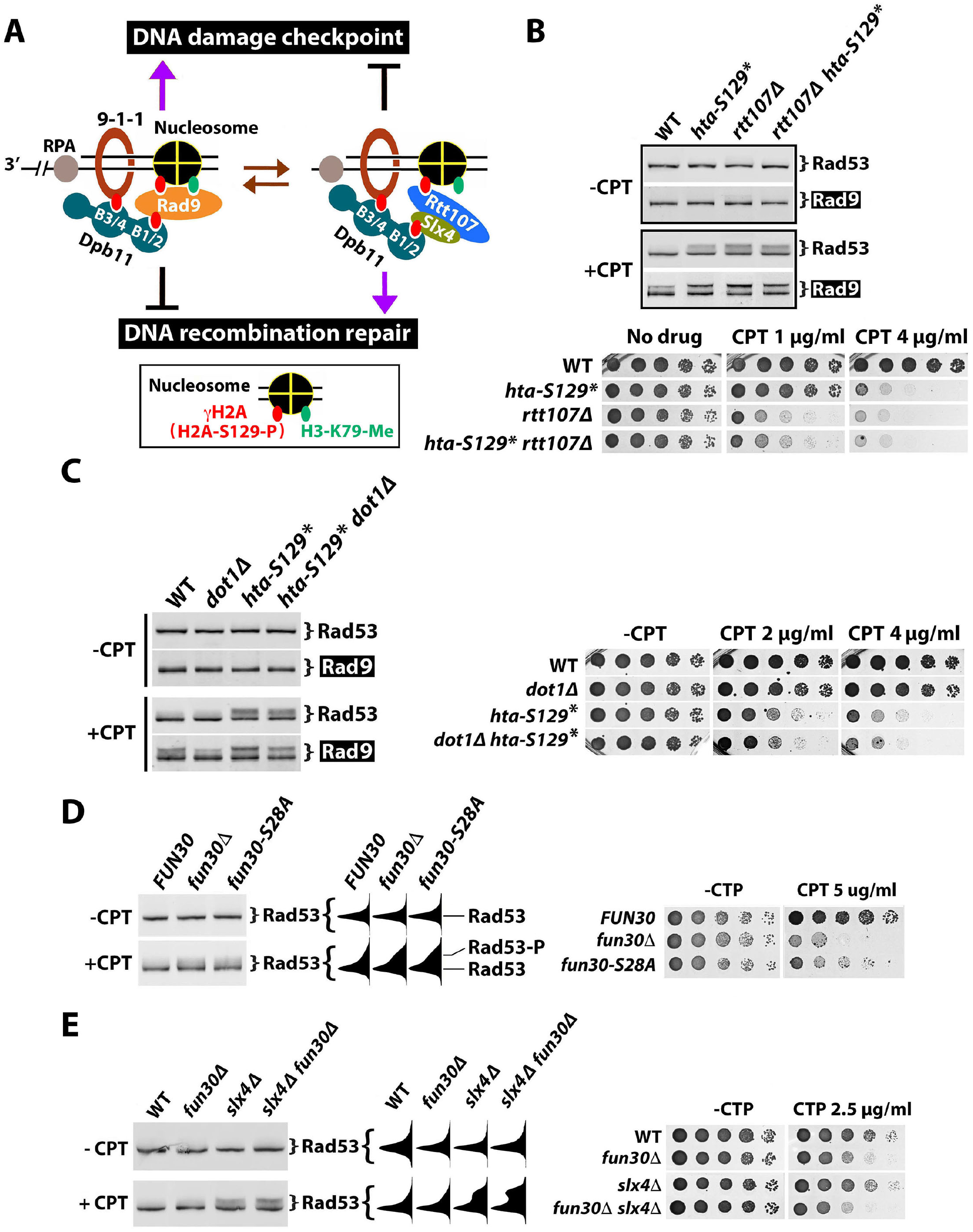
γH2A and Fun30 may down regulate DDC by hindering Rad9 recruitment to DNA lesions. (A) Model for negative regulation of DDC by γH2A and Fun30. See the text for descriptions. (B-E) Shown are results from Western blot analysis of Rad53 and Rad9-HA from indicated strains with or without CPT treatment as well as growth phenotypes of the strains on media with or without CTP. Strains tested are #43, 44, 51 and 52 (Table 1) in (B); #43, 44, 53 and 54 in (C); #37, 38 and 41 in (D); and #57-60 in (E). Note the bands and smears corresponding to Rad53 and Rad53-P in each blot in (D) and (E) were quantified/scanned using NIH ImageJ software. The profiles are displayed to the right of the blot image.

In the above model, the negative role of γH2A in DDC is mediated by its association with Rtt107 to damaged chromatin. Consistently, we showed that γH2A no longer negatively impact Rad9-P or Rda53-P in the absence of Rtt107 (Fig. 7B, top, compare *rtt107Δ* and *rtt107Δ hta-S129\** in +CPT panel). Note that the abundance of Rad53-P or Rad9-P in *rtt107Δ hta-S129\** mutant was slightly lower than that in *rtt107Δ* mutant (Fig. 7B, top, +CPT panel), indicating that γH2A makes a minor contribution to DDC signaling, which can be revealed when its prevailing negative function is eliminated (by removing Rtt107) (see Fig. S3D). We found that γH2A plays a smaller role in cellular resistance to CPT than Rtt107, and this role is lost in the absence of Rtt107 (Fig. 7B, bottom). In fact, we noticed that the *rtt107Δ hta-S129\** mutant was slightly more resistant to CPT than *rtt107Δ* mutant (Fig. 7B, bottom).

Unlike γH2A that can interact with both Rad9 and Rtt107, H3-K79-me is recognized by Rad9, but not Rtt107, and should therefore play a positive role in DDC. Consistently, H3-K79-me or Dot1 responsible for H3-K79 methylation has been shown to aid in DDC induced by phleomycin or ionizing radiation (IR) (37,61,81,82). We found here that *dot1Δ* reduces CPT-induced Rad9-P (Fig. 7C). However, CPT-induced Rad53-P level in *dot1Δ* cells was similar to, or only slightly lower than, that in WT (*DOT1*) cells (Fig. 7C). The lack of a robust effect of *dot1Δ* on Rad53-P might reflect the fact only limited Rad53-P was induced by CPT in WT cells to begin with (Fig. 7C). Deletion of *DOT1* moderately reduced CPT-induced Rda9-P and Rad53-P in *hta-S129\** cells (Fig. 7C), which is consistent with the notion that H3-K79-me helps recruit Rad9 independently of γH2A. Note that Rad9-P and Rad53-P in *hta-S129* dot1Δ* double mutant were more robust than those in *dot1Δ* and WT cells (Fig. 7C), suggesting that blocking Rad9 recruitment via association with chromatin marks (γH2A and H3-K79-me) does not markedly reduce DDC signaling. This is consistent with the notion that DDC signaling can proceed via Dpb11-mediated recruitment of Rad9 independently of the chromatin (γH2A and H3-K79-me) dependent pathway of Rad9 recruitment (61) (Fig. S3F). We showed although Dot1 contributes to DDC, it has little or no effect on CPT-resistance in the presence or absence of γH2A (Fig. 7C, right).

Fun30 was recently found to be phosphorylated at serine 28 (S28) by CDK in a cell cycle dependent fashion (62). Notably, phosphorylated Fun30 (Fun30-S28-P) interacts with B1/2 of Dpb11, the same domain that interacts with Rad9 and Slx4 phosphorylated by CDK (30,62,78). The fun30-S28A allele with S28 replaced by alanine fails to be phosphorylated and cannot interact with Dpb11 (30). Based on these finding, we posit that Fun30 inhibits Rad9 function by competing with it for binding Dpb11, thereby negatively regulating DDC. Consistent with this notion, we found that *fun30-S28A* mutation increased CPT-induced Rad53-P and made cells more sensitive to CPT, as *fun30Δ* did (Fig. 7D). Moreover, we showed that *fun30Δ* and *slx4Δ* had a synthetic effect on CPT-induced Rad53-P (Fig. 7E, left), indicating that Fun30 and Slx4 separately impact DDC. Slx4 and Fun30 also separately contribute to cellular resistance to CPT (Fig. 7E, right).

### 3.5. The effect of γH2A on checkpoint signaling is DNA damage-dependent

That γH2A exhibits a negative effect on CPT-induced G2/M checkpoint (Fig. 4 and 5) is not in line with previous studies suggesting positive or no functions of γH2A in DDC. For example, γH2A was found to contribute to G1 checkpoint but not G2/M checkpoint induced by IR or phleomycin (35,61,82). Moreover, γH2A was shown to play a minor role in intra-S checkpoint induced by MMS (35,61,82). Our work differs from these studies in the genotoxin used (CPT vs. IR, phleomycin or MMS) and genetic backgrounds of yeast cells used, which may influence the DDC responses examined (8). In an attempt to uncover the cause of the apparent discrepancies between our and other studies, we systematically tested the effects of *hta-S129\** on DDC in response to MMS and phleomycin in addition to CPT in the same genetic background of JKM139 (Table 1). Both Rad53-P and Rad9-P were measured as indicators of DDC signaling in exponentially growing cells. Effects of deleting known DDC regulators Slx4, Rtt107 and Sae2 on DDC were also monitored as controls.

As shown in Fig. 8A, *htaS-129A*, slx4Δ, rtt107Δ* and *sae2Δ* increased CPT-induced Rad53-P and Rad9-P to various degrees (compare panels CPT and WT), confirming that γH2A, Slx4, Rtt107 and Sae2 all play negative roles in DDC in response to CPT. MMS induced robust Rad53-P and Rda9-P in wild cells (Fig. 8A, MMS panel). The *hta-S129\** mutation increased MMS-induced Rad53-P, and to a lesser extent, Rad9-P (Fig. 8A, MMS panel), indicating that γH2A also negatively impact DDC signaling in response to MMS. Consistent with previous studies, *rtt107Δ, slx4Δ* and *sae2Δ* were all found to markedly increase MMS-induced Rad53-P (Fig. 8A, MMS panel) (55,56). We showed here that these mutations also enhanced Rad9-P in response to MMS (Fig. 8A, MMS panel).

**Fig. 8.**
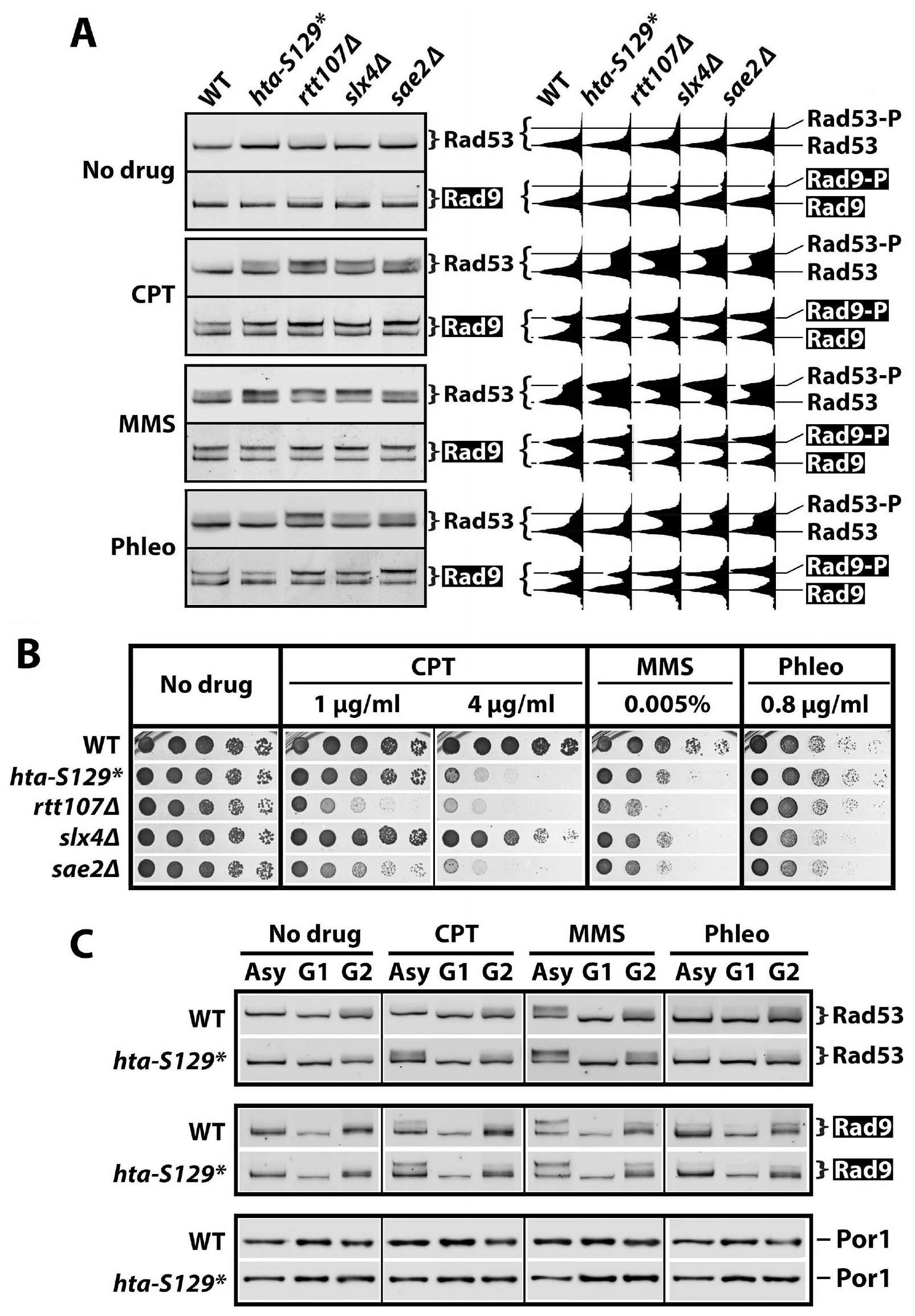
γH2A impacts DDC in a DNA damage-specific manner. (A) Western blot analyses of Rad53 and Rad9-HA (Rad9) from indicated strains (#43, 44, 51, 55 and 56 in Table 1) treated with CPT, MMS or phleomycin (Phleo), or mock treated (No drug). Exponentially growing cells of each strain were treated for 90 minutes with 5 μg/ml CPT, 0.01% MMS or 5 μg/ml phleomycin. Bands and smears corresponding to Rad53, Rad53-P, Rad9 and Rad9-P in the blots were quantified/scanned using NIH ImageJ software, and the profiles displayed to the right of the blot images. (B) Growth phenotypes of strains #43, 44, 51, 55 and 56 (Table 1) on media with or without the indicated genotoxins. (C) Impact of γH2A on checkpoint signaling in different phases of the cell cycle. An exponentially growing culture of WT (#49 in Table 1) or hta-S129* (#50) was divided into three aliquots, two of which were arrested in G1 and G2/M (designated G2 for simplicity) by α-factor and nocodazole, respectively, whereas the other one was not treated and remained asynchronous (Asy). Asy, G1 and G2 cultures were each divided into four aliquots, three of which were treated for 90 minutes with 5 μg/ml CPT, 0.01% MMS and 5 μg/ml phleomycin (Phleo), respectively, whereas the other was mock treated (No drug). Rad53, Rad9-HA (Rad9) and Por1 in each sample were examined by Western blotting.

Phleomycin induced a moderate level of Rad53-P and a relatively higher level of Rad9-P (Fig. 8A, Phleo panel). The *hta-S129\** mutation moderately reduced phleomycin-induced Rad53-P and Rad9-P (Fig. 8A, Phleo panel). On the other hand, the *rtt107Δ, slx4Δ* and *sae2Δ* mutations all increased both Rad53-P and Rad9-P in response to phleomycin (Fig. 8A, Phleo panel). Therefore, phleomycin-induced DDC signaling is partially dependent on γH2A, and is inhibited by Slx4/Rtt107 and Sae2.

Results from the above experiment revealed that γH2A plays a negative role in checkpoint signaling induced by CPT or MMS, but a positive role in checkpoint induced by phleomycin. This was corroborated by data obtained from independent experiments on the effects of *hta-S129A* on DDC signaling (Fig. S4A). Therefore, γH2A exhibits DNA damage-specific effects on DDC signaling, which is in contrast to Slx4, Rtt107 or Sae2 that dampens checkpoint induced by any DNA damage tested above.

We also examined how *hta-S129\** as well as *slx4Δ, rtt107Δ* and *sae2Δ* mutations affect cell survival in the presence of CPT, MMS or phleomycin. As shown in Fig. 8B, each mutation decreased cellular resistance to all three genotoxins to various degrees, except that *hta-S129\** and *rtt107Δ* did not have a significant effect on phleomycin resistance. The latter is interesting since *hta-S129\** and *rtt107Δ* had opposite effects on phleomycin-induced DDC signaling (Fig. 8A, Phleo panel). On the other hand, *slx4Δ* and *sae2Δ* mutants had similar levels of DDC signaling in the presence of CPT (Fig. 8A, CPT panel), but exhibited drastically different degrees of CPT-resistance (Fig. 8B). These results demonstrate again a lack of a straightforward correlation between the degree of DDC signaling and cell survival in the presence of DNA damage.

### 3.6. Impacts of γH2A on checkpoint signaling in different phases of the cell cycle

The above data revealing genotoxin-specific effects of γH2A were obtained from asynchronously growing cells, and thus did not reveal what checkpoint(s) (G1, intra-S or G2/M) was affected by γH2A. To address this question, we examined the levels of Rad53-P and Rad9-P in cells arrested in G1 or G2/M treated with CPT, MMS or phleomycin, and compared them to those in asynchronous cells. It is known that Rad53 and Rad9 are subject to phosphorylation in the absence of DNA damage by cyclin-dependent kinase CDK and Cdc5 (for Rad53 only) that is most prominent in G2/M phase (34,83–85). These DNA damage independent phospho-forms of Rad53 and Rad9 are referred to as Rad53-Pc and Rad9-Pc, respectively, and have faster mobility than the DNA damage-induced phospho-forms (referred to as Rad53-P and Rad9-P above) (33,83,85–87). Accordingly, we found Rad53-Pc and Rad9-Pc to be abundant in G2/M phase, less so in asynchronous cells, and hardly detectable in G1 phase (Fig. 8C and S4B, “No drug” panel). The *hat-S129\** mutation had little or no effect on Rad53-Pc and Rad9-Pc in asynchronous, G1 or G2/M cells (Fig. 8C and S4B, “No drug” panel), indicating that γH2A is not involved in DNA damage-independent phosphorylation of Rad53 and Rad9.

CPT treatment induced modest levels of Rad53-P and Rad9-P in asynchronous WT cells (Fig. 8C, compare CPT/Asy with No drug/Asy; also see Fig. S4C), which is consistent with results shown in Fig. 8A. CPT had little effect on Rad53-P and Rad9-P in G1 cells (Fig. 8C, compare CPT/G1 with No drug/G1). In G2/M cells, CPT increased Rad9-P, and to a lesser extent, Rad53-P (Fig. 8C, compare CPT/G2 with No drug/G2). The *hta-S129\** mutation significantly increased CPT induced Rad53-P and Rad9-P in asynchronous cells, but had little or no effect in G1 or G2/M cells (Fig. 8C, CPT panel). As a consequence, CPT-induced Rad53-P and Rad9-P in asynchronous *hta-S129\** cells were clearly more robust than those in G2 cells (Fig. 8C, CPT panel), suggesting that although CPT-induces a G2/M checkpoint (Fig. 3 and 5), activation of this checkpoint requires cells to traverse S phase. This is consistent with the notion that CPT-induced DNA lesions originate from encounters between Top1ccs and DNA replication forks in S phase, which triggers DDC signaling in G2/M. γH2A may function in S and/or G2/M phases to suppress DDC signaling induced by CPT.

MMS induced abundant Rad53-P and Rad9-P in asynchronous WT cells (Fig. 8C, MMS panel), as have been shown above (Fig. 8A). Lower levels of Rad53-P and Rad9-P were also observed in G2/M cells, but little was detected in G1 cells in the presence of MMS (Fig. 8C, MMS panel). MMS at the concentration used in this work (0.01%) would induce a salient G2/M arrest (see ref. 88; J.S. and X.B. data not shown). The fact MMS-induced Rad53-P and Rad9-P were much higher in asynchronous cells than G2/M cells (Fig. 8C, MMS panel) suggests that the activation of G2/M checkpoint in response to MMS also requires cell progression through S phase, similarly as CPT-induced G2/M checkpoint. This is likely because G2/M checkpoint induced by MMS is triggered by ssDNA at replication forks stalled by MMS mediated DNA methylation that is left behind after replication restart, and is subject to post-replication repair in G2/M phase (8). The *hta-S129\** mutation increased MMS-induced Rad53-P and Rad9-P in both asynchronous and G2/M cells (Fig. 8C, MMS panel), suggesting that γH2A can function in S and G2/M phases to suppress G2/M checkpoint in the presence of MMS.

Phleomycin induced modest levels of Rad53-P and Rad9-P in asynchronous and G1-arrested cells, and higher levels of Rad53-P and Rad9-P in cells arrested in G2/M (Fig. 8C and 4SC, Phleo panel). The *hat-S129\** mutation reduced Rad53-P and Rad9-P in G2/M cells (Fig. 8C and 4SC, Phleo panel). Note Rad53-P and Rad9-P were still readily detectable in *hat-S129\** cells in G2/M (Fig. 8C, Phleo panel), suggesting that γH2A plays only a partial role in the activation of G2/M checkpoint in response to phleomycin. We observed a moderate decrease in phleomycin-induced Rad53-P and Rad9-P in asynchronous or G1 cells as a result of *hta-S129\** mutation (Fig. 8C and S4C, Phleo panel).

The above results revealed that γH2A has a moderate negative effect on MMS-induced DDC signaling in G2/M phase, and a barely detectable negative effect on CPT-induced DDC in G2/M. On the other hand, γH2A has a positive effect on phleomycin-induced DDC signaling in G2/M as well as G1 albeit to a lesser extent. They also showed an S phase requirement for CPT-or MMS-induced G2/M checkpoint that is subject to suppression by γH2A.

## 4. Discussion

In this report we showed that Tel1 and γH2A have DNA damage-specific effects on DDC signaling. They negatively impact CPT-induced G2/M checkpoint independently of each other. γH2A also down regulates MMS-induced DDC signaling, but plays a positive role in DDC in response to phleomycin. Tel1 suppresses CPT-induced DDC in the same pathway as MRX possibly by processing/repairing Top1cc without yielding DDC-inducing DNA structures. γH2A may negatively impact DDC by serving as a platform for the recruitment of DDC inhibitor Slx4/Rtt107 to DNA lesions.

### 4.1. Tel1 negatively impacts DDC signaling in response to CPT

Although Tel1 is a homolog of mammalian ATM kinase, it does not play an essential role in DDC in yeast (3,4). For instance, *tel1Δ* slightly reduces, or does not affect, DDC signaling in response to DNA lesions induced by phleomycin, MMS, or endonucleases (5,68,71,72). It is intriguing that we found *tel1Δ* to enhance CPT-induced DDC signaling (Fig. 2A and 3), which was also reported recently by Menin et al. when this manuscript was being prepared (68). It is possible that the DNA lesion induced by CPT is distinct from those inflicted by the other abovementioned genotoxins, and that Tel1 plays a unique role in cellular response to this lesion. CPT is a topoisomerase I specific poison that traps/stabilizes Top1ccs that, upon collision with replication forks, can give rise to DSBs that induce DDC (66,89,90). Tel1 is known to participate in initial DSB resection and early DDC signaling in response to DSBs (9) (Fig. 1B). Tel1 phosphorylates histone H2A (to make γH2A) and Sae2 (Fig. 1B). However, we showed that the negative impact of Tel1 on DDC is not dependent on γH2A or Sae2 (Fig. 4A and S1D). We also showed that the effect of Tel1 on DDC signaling is independent of Fun30 chromatin remodeler that promotes long range DSB end resection (58) (Fig. 2A). These results suggest that the role of Tel1 in controlling CPT-induced DDC may be different from its function in cellular response to DSBs.

It has long been noticed that DDC signaling induced by CPT is relatively limited compared to that induced by other DNA damaging agents such as MMS (58,65). In fact, although Top1cc presents a potential obstacle to DNA replication, sublethal doses of CPT do not induce intra-S phase DDC or significantly delay S phase progression (Fig. 3) (58,65). There may be a mechanism(s) in the cell for preventing DNA replication forks from colliding with CPT-trapped Top1ccs on the DNA, thereby avoiding the formation of DDC-inducing DSBs. Consistent with this notion, there is evidence suggesting that CPT-trapped Top1cc can retard the incoming replication-fork and promote fork reversal (91). Fork stalling is likely due to the accumulation of positive supercoils in front of the fork as a consequence of the inhibition of Top1 activity by CPT (92). The reversed replication fork can be resolved by fork fusion or Top1cc repair, which does not generate a DSB or extended ssDNA region, thereby avoiding DDC activation (68,91). Consistent with this notion, we found that disrupting any of the 3 factors involved in the processing of Top1cc-like DNA-protein crosslinks (DPCs), Tdp1, Wss1 or MRX, leads to a significant increase in Rad53-P, similarly as *tel1Δ* (Fig. 6).

Tdp1, Wss1 and MRX are each able to promote Top1 removal from DNA (11,93–96). It is noteworthy that MRX functions in Top1 removal from DNA and cellular resistance to CPT are independent of its role in DSB repair (97). We found evidence suggesting that Tel1 acts in the same pathway as MRX in impacting CPT-induced DDC, and independently of Tdp1 and Wss1 (Fig. 6). That *mre11Δ, rad50Δ* and *xrs2Δ* are each epistatic to *tel1Δ* in increasing Rad53-P in the presence of CPT (Fig. 6) suggests that Tel1 acts upstream of MRX to suppress CPT-induced DDC signaling. We propose that Tel1 promotes the function of MRX in removing Top1 from DNA in the presence of CPT, thereby reducing the chance of collision between Top1cc and replication fork and DDC activation. Menin et al. reported that *tel1Δ* reduces the frequency of CPT-induced replication fork reversal and increases the abundance of long ssDNA regions at replication forks, both of which are suppressed by disrupting the nuclease activity of Mre11 (68). They therefore posited that Tel1 inhibits Mre11 mediated nucleolytic degradation of reversed replication fork that can lead to the formation of long ssDNA stretches and trigger DDC (68). It is possible that Tel1 suppresses CPT-induced DDC signaling both by positively regulating Top1 removal from DNA and by negatively regulating nucleolytic degradation of reversed replication fork.

### 4.2. DNA damage-specific impacts of γH2A on DDC

γH2A/γH2AX plays key roles in DNA damage response by serving as a docking site for multiple DDC or DNA repair factors including Rad9 and Slx4/Rtt107 (see Fig. 7A). Given the mutually exclusive nature of γH2A association with these factors, γH2A has the potential to positively or negatively regulate various processes involved in DNA damage response. For example, γH2A can serve dual, opposing functions in DDC activation, one is to promote DDC by recruiting Rad9 to DNA lesions, and the other is to inhibit DDC by recruiting Slx4/Rtt107 (56) (see Fig. 7A). The outcome of the competition between these two functions would determine whether γH2A has a net positive or net negative impact on DDC. As such, our finding that blocking γH2A increases DDC signaling induced by CPT or MMS (Fig. 8, A and C) likely reflects that γH2A-mediated recruitment of Slx4/Rtt107 outweighs Rad9 recruitment in the presence of CPT or MMS. On the other hand, Rad9 recruitment by γH2A seems to prevail over Slx4/Rtt107 recruitment in response to phleomycin or IR, as the lack of γH2A reduces phleomycin-or IR-induced DDC (35,61) (see Fig. 8, A and C).

That γH2A differentially impacts DDC in response to CPT, MMS, phleomycin or IR can be explained by assuming that the result of the competition between Rad9 and Slx4/Rtt107 for binding γH2A at a DNA lesion is dependent on the nature/type of the lesion. Both CPT and MMS trigger DDC by inducing replicative stress in S phase, whereas phleomycin and IR can induce DSBs and DDC in any phase of the cell cycle (see Fig. 8C). It is tempting to propose that a DNA replication stress-induced DNA lesion presents a more favorable context for Slx4/Rtt107-γH2A interaction than Rad9-γH2A interaction, whereas a DSB induced by phleomycin or IR is more favorable for Rad9-γH2A interaction.

It is noteworthy that although CPT and MMS each trigger a G2/M checkpoint (Fig. 3 and 5), checkpoint activation requires the host cell to traverse S phase (58,65) (Fig. 8C). CPT-trapped Top1cc or MMS-mediated DNA methylation stalls DNA replication in S phase, which has to be resolved to allow the completion of DNA replication. Mechanisms for the restart of DNA replication may involve the generation of altered fork structures (as fork restart intermediates) harboring ssDNA gaps that can trigger DDC (98). We imagine that these distorted fork structures are different from resected (simple) DSBs induced by phleomycin or IR in influencing γH2A interaction with Rad9 and/or Slx4/Rtt107. Specifically, we posit that Slx4/Rtt107 outcompetes Rad9 for binding γH2A within the fork restart intermediates induced by CPT or MMS, whereas Rad9 is favored for binding γH2A at resected DSBs in the presence of phleomycin or IR. Consequentially, γH2A plays a net negative role in DDC signaling in response to CPT or MMS, but a positive role in DDC triggered by phleomycin or IR.

### 4.3. Relationship between DDC signaling and cell survival in the presence of genotoxins

Cells defective in proper DDC signaling are usually sensitive to genotoxins (1). However, our survey of DDC signaling and CPT-sensitivities of the series of mutants described in this work revealed a lack of a causative link between the level of DDC signaling and the degree of CPT-resistance, which is line with prior observations (99–101). This could be because efficient cellular resistance to a genotoxin requires an “optimal” level of DDC signaling. Alternatively, or in addition, genotoxin resistance may reflect aggregated effects of the genotoxin on different aspects of cellular response to DNA damage. Given that many factors involved in DNA damage response have multiple functions that are not restricted to DDC, deletion of a particular factor may have effects on not only DDC, but also DNA replication and/or DNA repair in the presence of a genotoxin. The combination of these effects likely determines the ability of the mutant to withstand DNA damage induced by the genotoxin. Separation-of-function mutations of DDC factors would be particularly useful for examining how these factors contribute to DCC signaling and genotoxin resistance.

## Acknowledgement

We thank Ms. Carina Wong and Dr. Anhui Wei for assistance. We thank Drs Thomas Petes (Duke University), Stephen Kron (University of Chicago), James Haber (Brandeis University), Grzegorz Ira (Baylor College of Medicine) and Elizabeth Grayhack (University of Rochester) for gifts of yeast strains. This work was supported by NSF grant MCB-1158008 to X.B.

---

**Fig. S1.**
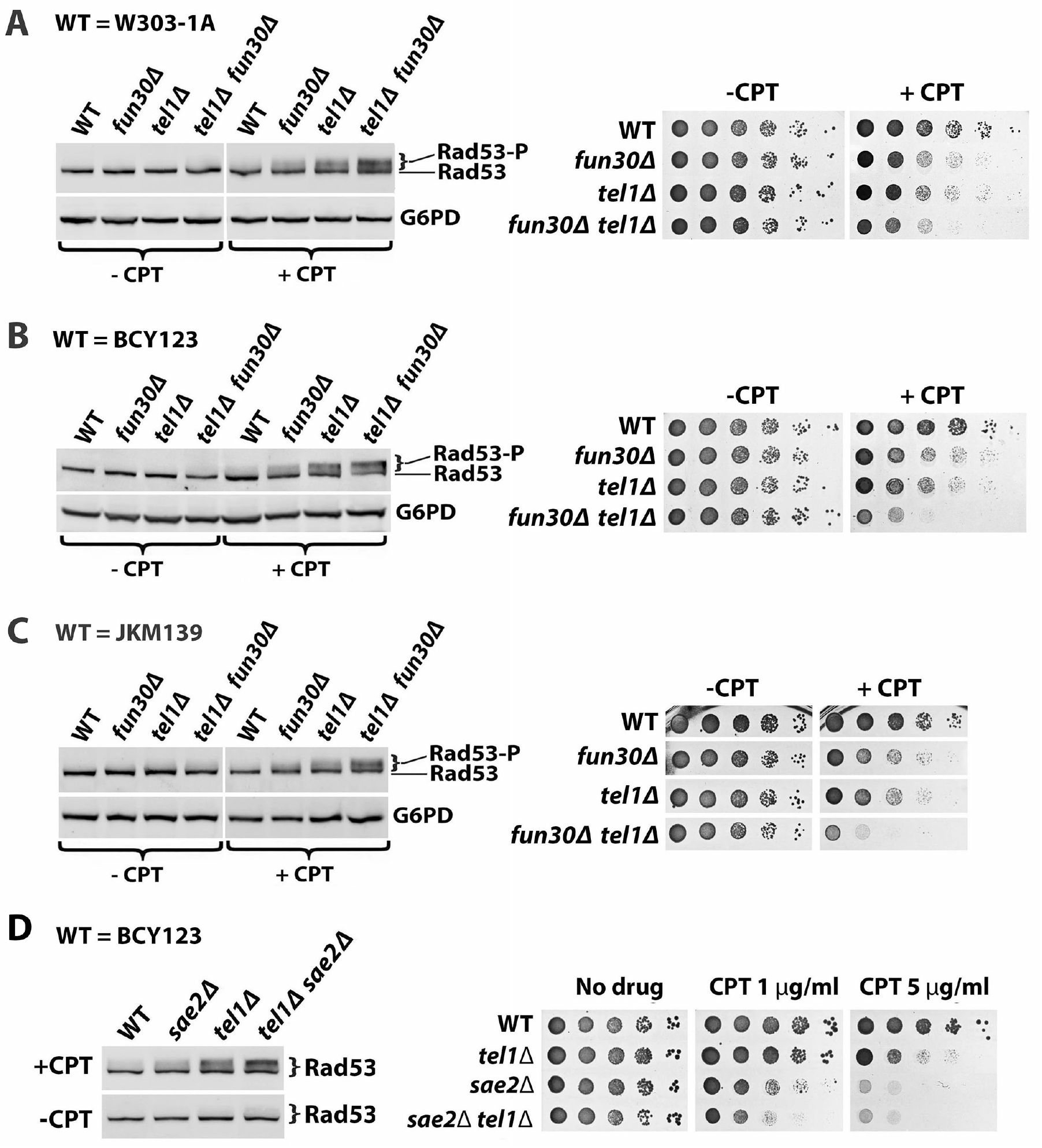
Functional relationships of Tel1 with Fun30 and Sae2 in affecting DDC signaling and CPT-resistance. Shown are data from Western blot analysis of Rad53 and G6PD from indicated strains with or without CPT treatment (5 μg/ml for 90 minutes) as well as growth phenotypes of the strains on media with or without CTP. (A) W301-1A and its derivatives (#1-4 in Table 1) were tested. (B) BCY123 and its derivatives (#14-17 in Table 1) were tested. (C) JKM139 and its derivatives (#37-40 in Table 1) were tested. (D) BCY123 and its derivatives (#14, 16, 28 and 29 in Table 1) were tested.

**Fig. S2.**
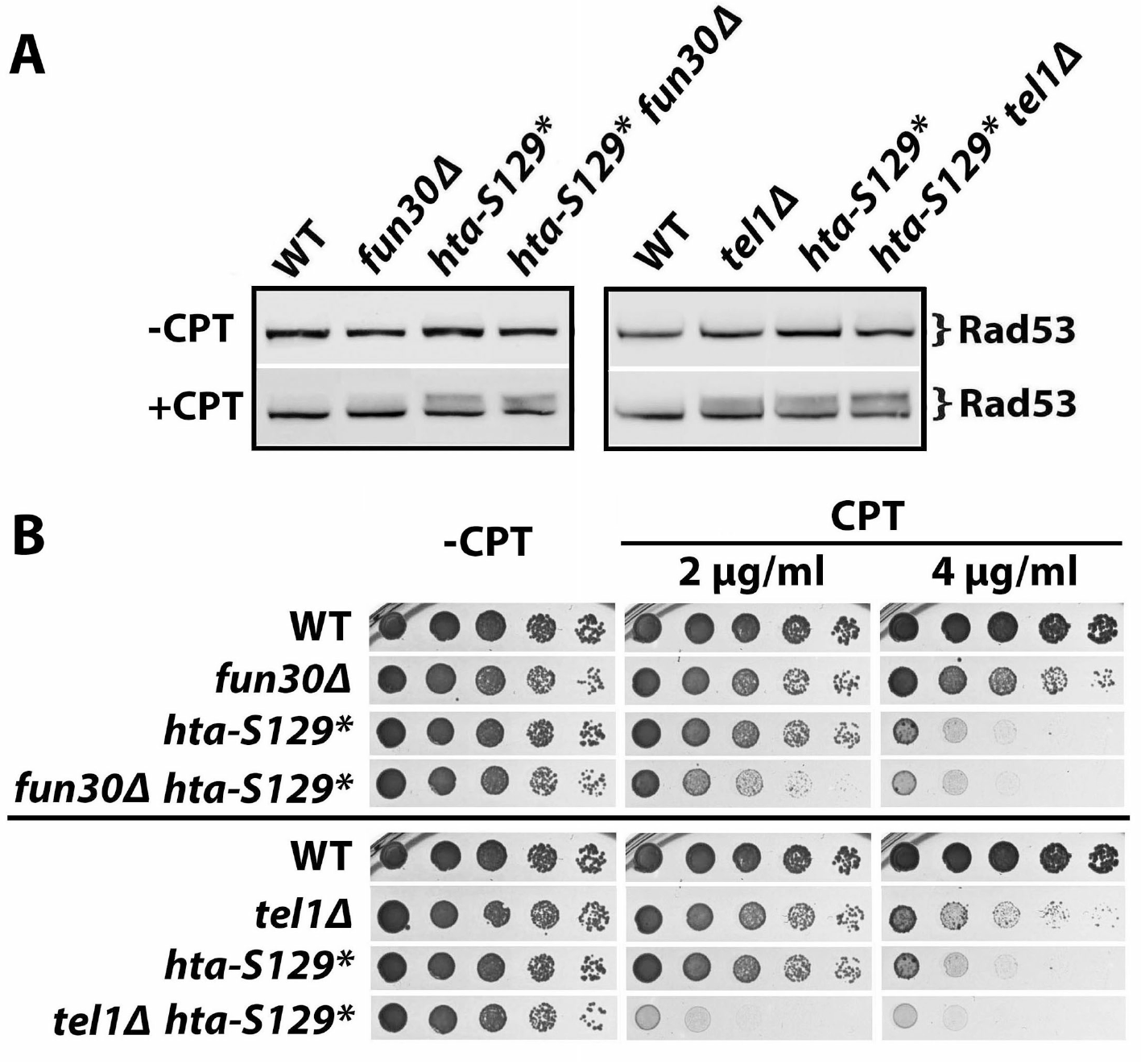
Effects of *hta-S129\** mutation on CPT-induced Rad53-P and CPT-resistance. (A) Western blot analysis of Rad53 from indicated strains (#43-48 in Table 1) with (+CPT) or without (-CPT) CTP treatment. (B) Growth phenotypes of indicated strains (#43-48 in Table 1) on media with or without CTP.

**Fig. S3.**
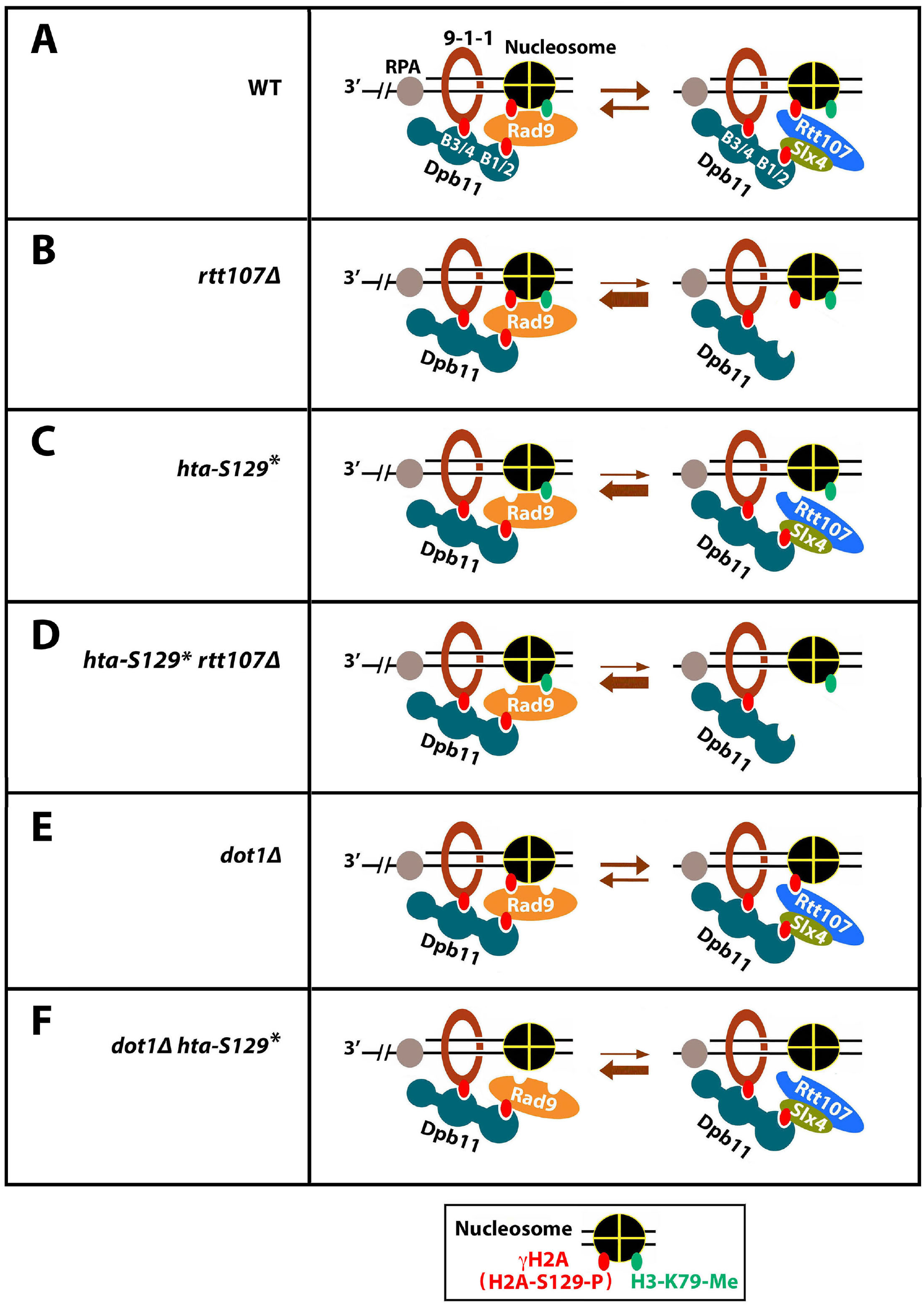
Illustration of the competition between Rad9 and Slx4/Rtt107 for binding γH2A and Dpb11 in various genetic backgrounds in response to DNA damage. See the text for descriptions. The genetic background considered here are (A) Wild type, (B) *rtt107Δ*, (C) *hta-S129\**, (D) *hta-S129* rtt107Δ*, (E) *dot1Δ*, and (F) *dot1Δ hta-S129*.*

**Fig. S4.**
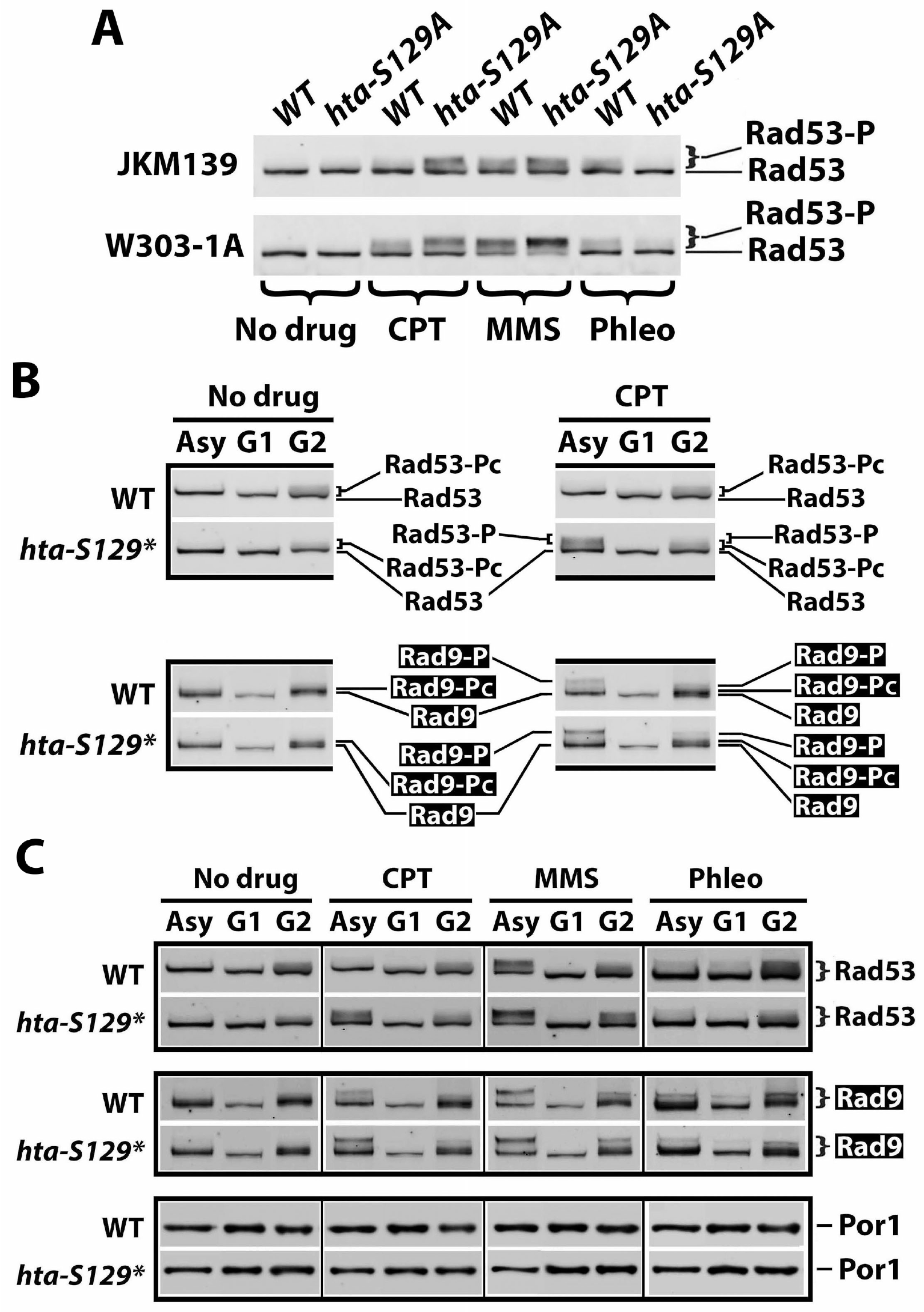
The effect of γH2A on DDC is DNA damage-specific. (A) Western analysis of Rad53 from JKM139 and R726 (JKM139, *hta-S129A)* as well as W303-1A and SKY2939 (W303-1A, *hta-S129A)* treated with 5 μg/ml CPT, 0.01% MMS or 5 μg/ml phleomycin, or mock treated (No drug). (B) Part of Fig. 8C with bands/smears corresponding to Rad53, Rad53-P, Rad53-Pc, Rad9, Rad9-P, and Rad9-Pc indicated. (C) A darker image of Fig. 8C.

